# Protective mechanisms against Alzheimer’s Disease in APOE3-Christchurch homozygous astrocytes

**DOI:** 10.1101/2025.01.21.634115

**Authors:** Xinran Tian, Erica Keane Rivera, Arina A. Nikitina, Inyoung Hwang, Jennifer A. Smith, Youngsuh J. Cho, Julia Sala-Jarque, Austin DuBose, Clare Andriola, Toby Gollan-Myers, Kenneth S. Kosik

## Abstract

The APOE3-Christchurch (APOE3-Ch) variant has been linked to reduced Alzheimer’s Disease (AD) risk, but its protective mechanisms remain unclear. This study explores the neuroprotective phenotype of APOE3-Ch astrocytes, focusing on lipid metabolism and tau processing. APOE3-Ch astrocytes demonstrate enhanced tau oligomer uptake via HSPG- and LRP1-mediated pathways, facilitated by elevated HSPG expression, and achieve superior tau degradation through lysosomal pathways and proteasomal pathways, in contrast to wild-type astrocytes, which primarily use proteasomal mechanisms. Transcriptomic analysis reveals upregulation of genes involved in endocytosis and cell projection assembly, explaining enhanced tau uptake and clearance in APOE3-Ch astrocytes. Lipidomic profiling identifies reduced levels of pathological lipids such as ceramides and gamma-linolenic acid (GLA), potentially mitigating neuroinflammation. These findings provide insight into the protective mechanisms of APOE3-Ch astrocytes and underscore their potential as therapeutic targets for tauopathy and neurodegeneration in AD.

**Teaser:** APOE3-Christchurch astrocytes enhance tau clearance and mitigate neurotoxic lipid accumulation, unveiling protective mechanisms against Alzheimer’s.

## Introduction

Astrocytes are critical for maintaining central nervous system (CNS) homeostasis, playing essential roles in lipid metabolism and the clearance of toxic protein aggregates. In addition to regulating extracellular ion and neurotransmitter levels, astrocytes metabolize excess lipids from neurons during periods of heightened neuronal activity, preventing lipotoxicity and supporting neuronal function (*1*, *2*). Central to these processes is apolipoprotein E (APOE), a lipid-binding protein expressed by astrocytes, which facilitates cholesterol transport, extracellular amyloid-beta (Aβ) clearance, synaptic pruning, and blood-brain barrier (BBB) integrity (*2*). The functional diversity of APOE isoforms profoundly impacts astrocyte biology (*3–5*).

While extensive research has focused on the common APOE isoforms (APOE2, APOE3, and APOE4), much less is known about rare variants such as APOE3-Christchurch (APOE3-Ch). APOE3-Ch has been associated with reduced AD risk and neuroprotection in individuals carrying high-risk genetic mutations for AD (*6–10*). However, the underlying mechanisms of protection remain poorly understood.

In addition to their role in lipid metabolism, astrocytes are increasingly recognized as active participants in tau clearance, a process critical for limiting the spread of pathological tau aggregates in AD and related tauopathies (*11–13*). Pathological tau, characterized by hyperphosphorylation and aggregation into neurofibrillary tangles, disrupts synaptic integrity and neuronal connectivity, driving neurodegeneration and cognitive decline (*14*, *15*). The mechanisms of tau uptake in astrocytes remain underexplored. The efficiency of tau processing is influenced by APOE isoforms, with APOE4 impairing tau uptake and trafficking, potentially exacerbating tau pathology (*16*).

To explore the protective mechanisms conferred by APOE3-Ch, this study investigates the lipid metabolism and tau processing dynamics of APOE3-Ch astrocytes compared to wild-type (WT) astrocytes. Using iPSC-derived astrocytes, we combined lipidomic profiling, transcriptomic analysis, and functional assays to characterize their phenotypic and functional responses. By delineating the unique lipid and tau regulatory mechanisms of APOE3-Ch astrocytes, this study offers novel insights into the molecular underpinnings of astrocyte-mediated neuroprotection. These findings highlight the potential of targeting astrocytic pathways as a therapeutic strategy for AD and related tauopathies.

## Results

### Characterization of iPSC-Derived APOE3-Christchurch and Wild-Type Astrocytes

To examine the phenotypic differences caused by the APOE3-Ch variant in astrocytes, We differentiated human induced pluripotent stem cells (hiPSCs) homozygous for either APOE3 or APOE3-Ch into astrocytes using a dual-SMAD inhibition-directed differentiation protocol (*17*). The parental iPSC line carrying the homozygous APOE3 is referred to as “P” or wild type (WT) in this study (Fig.1A). This line was subsequently modified using CRISPR to introduce the homozygous APOE3-Ch mutation. For this study, we selected two CRISPR-edited APOE3-Ch clones, labeled “1262” and “1264” (Fig.1A).

**Fig.1:**
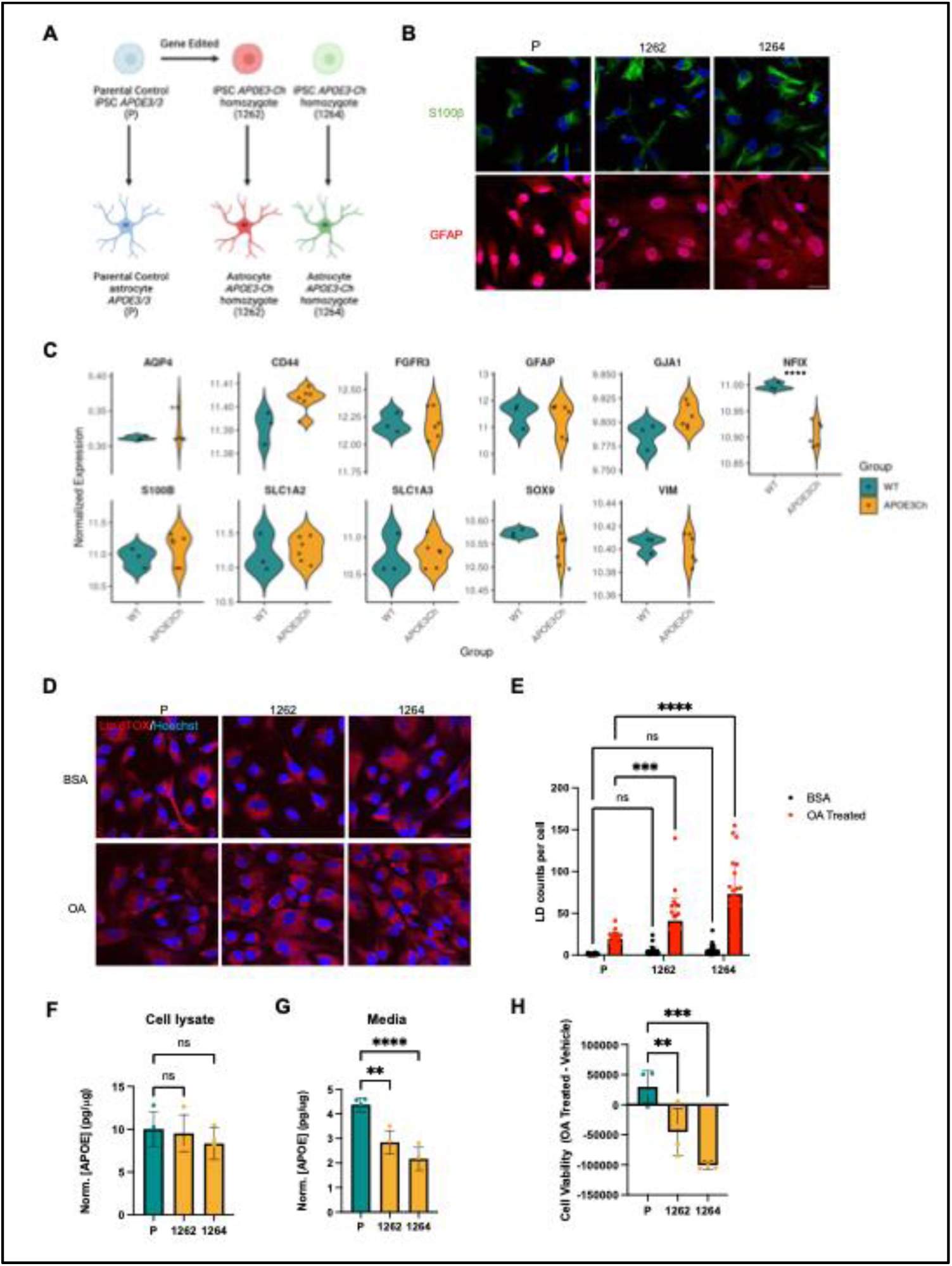
Characterization of iPSC-derived astrocytes and their lipid homeostasis. **(A)** Schematic representation of the experimental model. Gene-edited induced pluripotent stem cell (iPSC)-derived astrocytes were generated from parental control iPSCs expressing APOE3/3 (P) and from iPSCs homozygous for APOE3-Ch (1262 and 1264). **(B)** Immunofluorescence staining for the astrocytic markers S100β (green) and GFAP (red) in parental (P), 1262, and 1264 astrocytes. Nuclei are counterstained with Hoechst (blue). Scale bar represents 20 μm. **(C)** Violin plots representing normalized expression levels of astrocyte marker genes in APOE3-Ch (n=6) and WT astrocytes (n=3). P-values were calculated using a Wald test for differential expression between groups (WT vs APOE3Ch). Multiple testing correction was performed using the Benjamini-Hochberg method to control the false discovery rate (FDR). Genes with an adjusted p-value < 0.05 were considered statistically significant. **(D)** Lipid droplet (LD) staining using LipidTOX (red) and nuclei staining with Hoechst (blue) in P, 1262, and 1264 astrocytes following treatment with BSA (control) or oleic acid (OA). Scale bar represents 20 μm. **(E)** Quantification of lipid droplet counts per cell in P, 1262, and 1264 astrocytes under BSA or OA treatment conditions (n= 25 FOVs taken from 5 biological replicates, 5 FOVs per biological replicate). Significance was determined by two-way ANOVA with Dunnett’s multiple comparison tests. **(F-G)** APOE protein levels normalized to total protein (pg/μg) in cell lysates (f) and media (g) of P, 1262, and 1264 astrocytes (n=3). Significance was determined by one-way ANOVA with Dunnett’s multiple comparison tests **(H)** Cell viability following oleic acid (OA) treatment relative to vehicle control in P, 1262, and 1264 astrocytes (n=3). All data are expressed as mean ± s.d. with individual data points shown. The results designated as “ns” are not significant; **p<0.01; ***p<0.001;****p<0.0001

Following differentiation, we confirmed the expression of mature astrocyte markers S100B and GFAP through immunocytochemistry in both WT and APOE3-Ch astrocytes, indicating successful differentiation (Fig.1B). RNA sequencing further validated that these iPSC-derived astrocytes express high levels of key astrocyte markers, such as FGFR3, GFAP, S100B, and SLC1A2 (Fig.1C). Notably, however, these astrocytes exhibited low expression of AQP4, a marker of astrocyte maturation critical for water transport and potassium ion buffering. Reduced AQP4 expression may alter water and ion homeostasis in these cells.

Compared to WT astrocytes, APOE3-Ch astrocytes displayed significantly lower expression of the transcription factor NFIX (Fig.1C). NFIX plays a critical role in astrocyte differentiation and maturation. Its reduced levels in APOE3-Ch astrocytes suggest that these cells may be less mature than the control line, indicating that the APOE3-Ch variant may impede normal astrocyte maturation. Further investigation into how this variant affects astrocyte development and function is warranted.

### APOE3-Christchurch Astrocytes Exhibit Lipid Dyshomeostasis

APOE is a key component of lipoprotein particles such as chylomicrons, very-low-density lipoproteins (VLDL), and high-density lipoproteins (HDL). In the central nervous system, APOE facilitates the transport of cholesterol and fatty acids between cells, maintaining lipid homeostasis. This neuron-astrocyte metabolic coupling, mediated by APOE, plays an essential role in preventing activity-induced lipotoxicity in neurons (*1*). Clinical studies have suggested that the Christchurch variant may predispose individuals to hyperlipoproteinemia type III (HLPP3), characterized by elevated blood cholesterol and triglyceride levels, early-onset atherosclerosis, and heart disease (*18*).

To explore whether APOE3-Ch astrocytes exhibit lipid metabolism dysfunction, we quantified lipid droplet formation in both untreated astrocytes and those treated with oleic acid. Lipid droplets form to buffer excess intracellular lipids. In the absence of oleic acid treatment, APOE3-Ch astrocytes exhibited a trend toward increased lipid droplet production, though not significantly (Fig.1, D and E). However, upon oleic acid treatment, APOE3-Ch astrocytes produced significantly more lipid droplets than WT astrocytes, suggesting a reduced ability to secrete excess lipids. To assess whether this dysregulation is related to APOE secretion, we measured both intracellular and extracellular APOE levels using an enzyme-linked immunosorbent assay (ELISA). While intracellular APOE levels did not differ significantly between APOE3-Ch and WT astrocytes (Fig.1F), extracellular APOE levels were significantly lower in APOE3-Ch astrocytes (Fig.1G). This finding indicates a diminished capacity for APOE secretion in APOE3-Ch astrocytes, potentially leading to the intracellular accumulation of lipid droplets. This lipid dyshomeostasis may induce lipotoxicity and disrupt metabolic coupling between astrocytes and neurons. To further investigate whether this impaired lipid buffering induces cellular toxicity, we assessed astrocyte viability with an ATP-based cell viability assay after oleic acid treatment. APOE3-Ch astrocytes displayed decreased viability in the presence of excess lipids compared to WT astrocytes (Fig.1H). These results shed light on how the APOE3-Ch variant may contribute to lipid-related pathology, though additional studies are needed to elucidate the mechanisms underlying reduced APOE export and its effects on lipid metabolism.

### Lipidomic Analysis Reveals Reduced Ceramide Levels in APOE3Ch Astrocytes

Astrocytes are the main lipid producers in the brain. To explore the impact of the APOE3-Ch variant on astrocytic lipid metabolism, we performed targeted lipidomic profiling on both the cell pellet fractions and media fractions of astrocytes, aiming to uncover differences in the content of intracellular versus secreted lipids.

The intracellular neutral lipid panel revealed that cholesterol esters (CEs), ceramides (Cer), diacylglycerols (DAGs), and triacylglycerols (TAGs) were the more abundant lipid classes overall, while dihydroceramides (DCER) and monoacylglycerols (MG) exhibited lower abundances (Fig.2A). Notably, no significant differences were observed in the total abundance of these lipid species between APOE3-Ch astrocytes and wild-type (WT) astrocytes.

**Fig.2:**
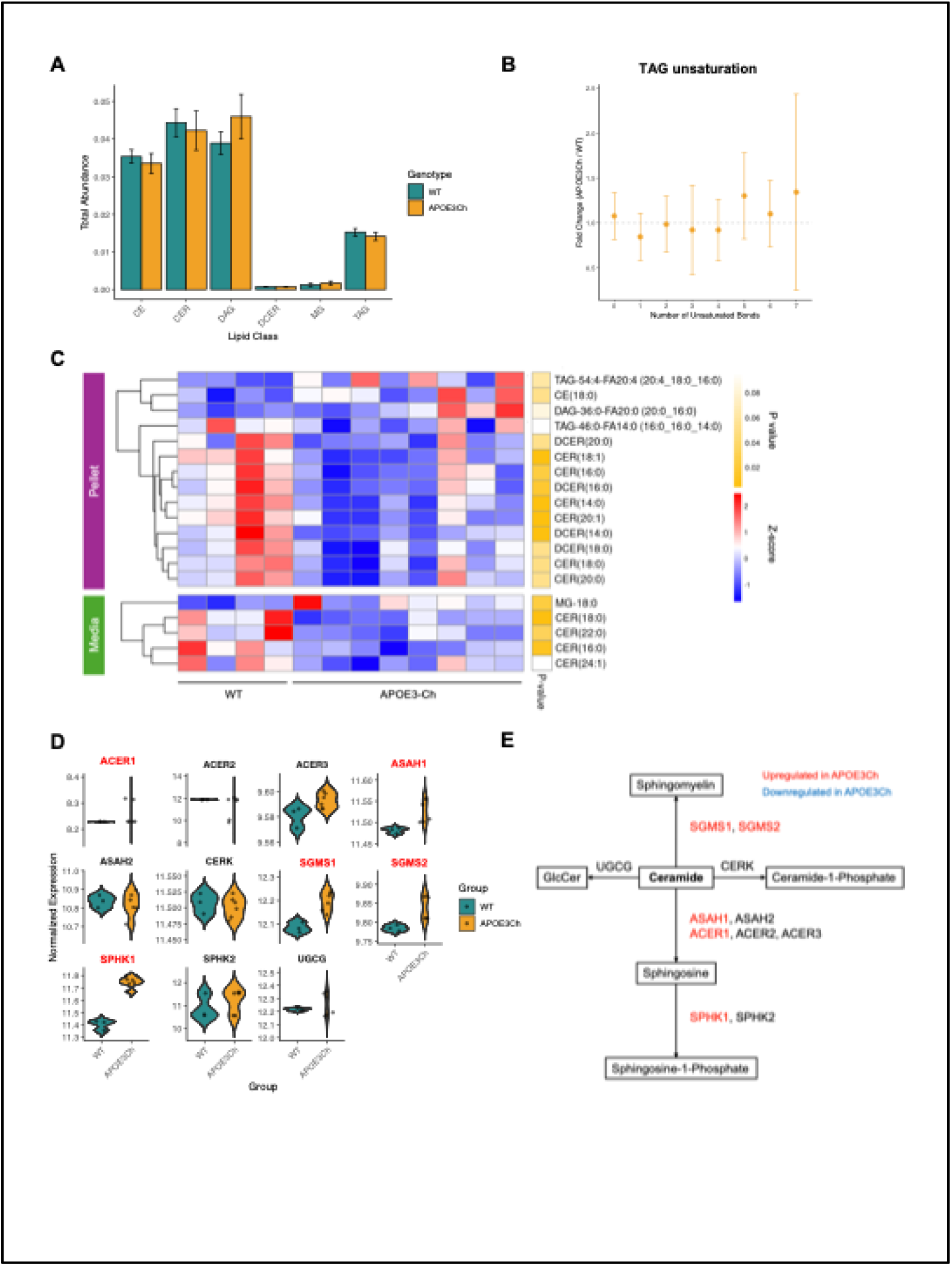
The neutral lipid profile suggests reduced ceramide levels in APOE3Ch astrocytes. **(A)** Total lipid abundance for each lipid species detected in the neutral lipid panel, comparing APOE3Ch (n=8) and WT astrocytes (n=4). Lipid abundance is represented as the sum of normalized peak area for each lipid class. Error bars denote the standard error of the mean (SEM). **(B)** Fold change in triacylglycerol (TAG) unsaturation levels between APOE3-Ch and WT astrocytes. The error bars represent the SEM. **(C)** Heatmap showing differentially abundant lipids in the pellets (intracellular) and media (extracellular) of APOE3-Ch astrocytes compared to WT astrocytes. Lipid expression is represented as a z-score, calculated by subtracting the mean lipid quantity across all samples from the observed quantity and dividing by the standard deviation. Increased and decreased lipids are color-coded. P-values were calculated using a linear model with empirical Bayes moderation of standard errors. **(D)** Violin plots representing normalized expression levels of genes involved in ceramide catabolism in APOE3-Ch (n=6) and WT astrocytes (n=3). P-values were calculated using a Wald test for differential expression between groups (WT vs APOE3-Ch). Multiple testing correction was performed using the Benjamini-Hochberg method to control the false discovery rate (FDR). Genes with an adjusted p-value < 0.05 were considered statistically significant. Significantly upregulated genes were highlighted in red. **(E)** Schematic representation of the ceramide catabolism pathway. Genes significantly upregulated in APOE3Ch astrocytes are highlighted in red, while downregulated genes are shown in blue. Downregulated genes were not found.

Given previous findings of elevated unsaturated TAGs in APOE4 astrocytes, we examined whether APOE3-Ch astrocytes displayed a similar trend. However, lipidomics analysis of the intracellular neutral lipid profile revealed that APOE3-Ch astrocytes exhibited levels of unsaturated TAGs comparable to those observed in WT astrocytes. (Fig.2B).

We further analyzed differentially abundant lipid species in both cell pellets and media (Fig.2C). In cell pellets, several ceramides and dihydroceramides were significantly reduced, including Cer (14:0), Cer (16:0), Cer (18:0), Cer (18:1), Cer (20:0), Cer (20:1), DCER (14:0), DCER (16:0), and DCER (18:0). Similarly, in the media, Cer (16:0), Cer (18:0), and Cer (22:0) were also significantly reduced. Ceramides are bioactive sphingolipids involved in cell signaling, apoptosis, and membrane structure, playing a critical role in neurodegenerative diseases like AD (*19*, *20*). Elevated ceramide levels have been associated with AD pathology, including promoting amyloid-beta accumulation, mitochondrial dysfunction, and inflammation (*21*, *22*). Specifically, increased levels of Cer16, Cer18, Cer20 and Cer24 are found in the brains from patients with AD (*23*). Therefore, lower levels of aforementioned ceramides, as observed in APOE3-Ch astrocytes, may mitigate these pathological processes, offering a protective effect against AD progression.

To investigate the underlying mechanisms for ceramide reduction, we assessed the expression of genes involved in ceramide catabolism from the RNAseq data (Fig.2, D and E). Several key enzymes, including ACER1, ASAH1, SGMS1, SGMS2, and SPHK1, were significantly upregulated in APOE3Ch astrocytes. Specifically, SGMS1 and SGMS2, which convert ceramides to sphingomyelin, were upregulated. ASAH1 and ACER1 are responsible for converting ceramides to sphingosine, while SPHK1 phosphorylates sphingosine to form sphingosine-1-phosphate (S1P), potentially further reducing ceramide levels by diverting sphingosine into the S1P pathway.

We also examined the polar lipid profile (fig.S1) and fatty acid composition (fig.S2) in APOE3Ch astrocytes. In the polar lipid panel, intracellularly, the most abundant lipids in both WT and APOE3-Ch astrocytes were phosphatidylcholine (PC), phosphatidylethanolamine (PE), phosphatidylinositol (PI), and phosphatidylserine (PS), with no significant differences in total abundance between genotypes (fig.S1A). However, several PE species were notably decreased, while several PS species were increased in APOE3-Ch astrocytes (fig.S1, B and C). In the media, differentially abundant polar lipids were scarce, though we observed significant downregulation of PE (14:0_16:0), PE (14:0_20:1), PC (14:0_14:0), and LPC (18:2) (fig.S1, D and E).

The fatty acid panel also revealed minimal differences in lipid species between APOE3-Ch and WT astrocytes. However, intracellularly, gamma-linolenic acid (GLA) and lauric acid were significantly downregulated (fig.S2, A and C) in APOE3-Ch astrocytes. Extracellularly, caproic acid was more abundantly exported in APOE3-Ch astrocytes compared to WT (fig.S2, B and C). Notably, dihomo-gamma-linolenic acid (DGLA), a metabolite of the polyunsaturated fatty acid GLA, has been implicated in inducing ferroptosis-mediated neurodegeneration, particularly in dopaminergic neurons (*24*). While the role of GLA in astrocytes remains unclear, we hypothesize that the downregulation of GLA in APOE3-Ch astrocytes may help prevent ferroptosis-mediated pathology, contributing to the protective mechanisms associated with this genotype.

Together, these findings suggest a genotype-specific remodeling of lipid metabolism that may contribute to the distinct functional properties of APOE3-Ch astrocytes.

### APOE3-Christchurch Astrocytes Internalize More Tau Oligomers

In addition to facilitating lipid metabolism, astrocytes also play a critical role in maintaining synaptic homeostasis by clearing harmful materials, including protein oligomers such as tau. Previous studies have demonstrated that astrocytes can internalize both tau monomers and higher-molecular-weight tau species (*13*, *25*). To investigate how the APOE3-Ch variant affects tau uptake, we prepared soluble 2N4R-P301L tau oligomers (fig.S3A), labeled them with Alexa Fluor 647 (AF647), and assessed tau uptake in APOE3-Ch and WT astrocytes via flow cytometry (Fig.3A). Our results revealed a significantly increased median AF647 intensity in APOE3-Ch astrocytes, indicating enhanced tau uptake compared to WT astrocytes (Fig.3, B and C). This difference became more pronounced with increasing tau concentrations and longer incubation times (fig.S3, B and C).

**Fig.3:**
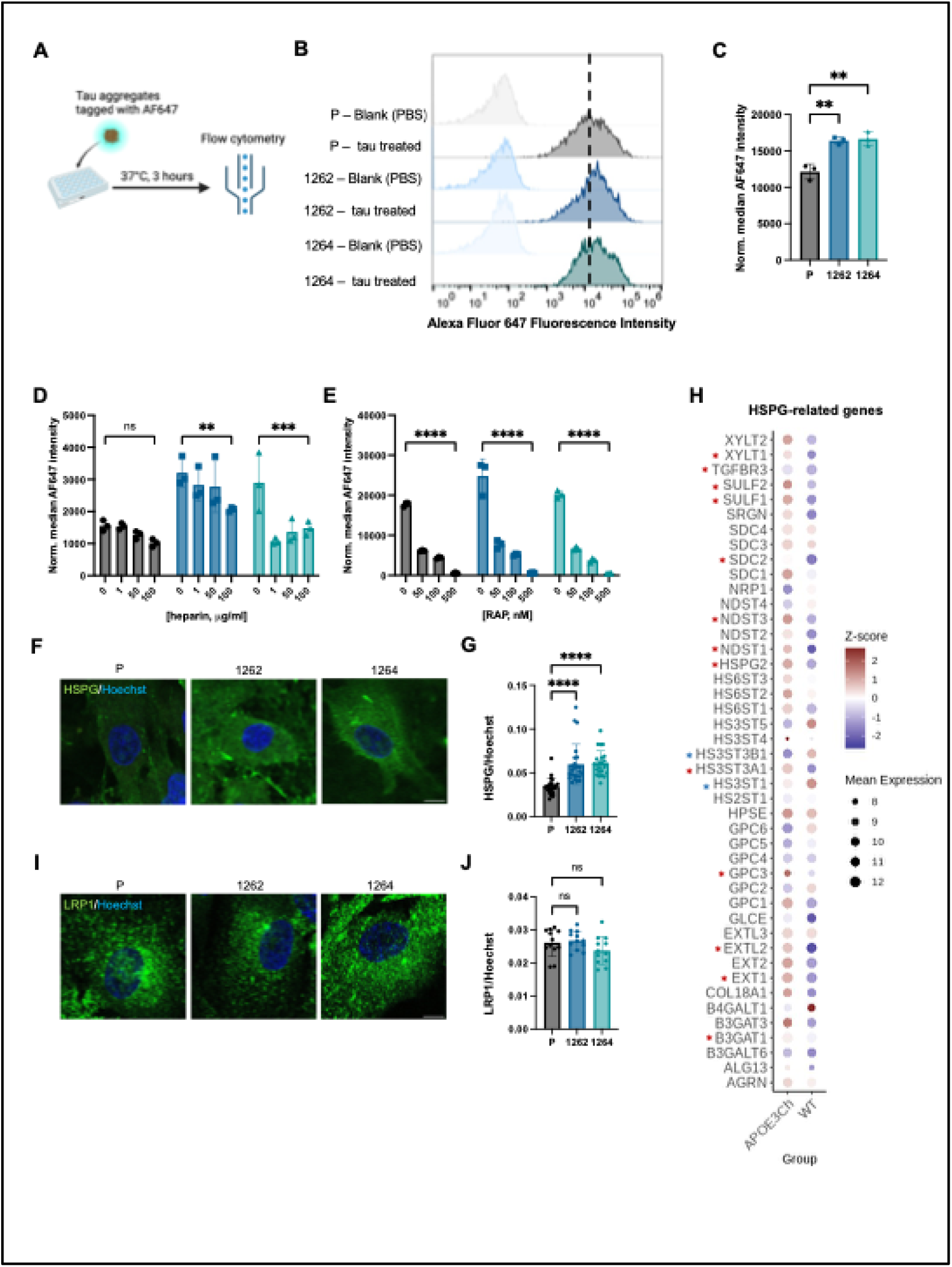
APOE3-Ch astrocytes take up more Tau aggregates. **(A)** Schematic of the tau uptake assay in iPSC-derived astrocytes. **(B)** Representative overlaid histograms demonstrating the shift in fluorescence of cells with AF647 labeled tau (2N4R-P301L) aggregates (50nM, 3h) compared to cells treated with PBS. The dotted line represents the median AF647 intensity in the WT astrocytes treated with Tau. **(C)** Measurement of tau uptake by flow cytometry normalized to cells treated with PBS (n=3, 10000 cells each). **(D)** Measurement of tau uptake in cells treated with Tau (50nM, 3h) and various concentrations of heparin normalized to cells treated with PBS (n=3, 10000 cells each). Statistical significance is determined by two-way ANOVA with uncorrected Fisher’s LSD multiple comparison tests. Displayed is the multiple comparison of cells treated with 100 ug/ml heparin against cells with no treatment for each cell line. **(E)** Measurement of tau uptake in cells treated with Tau (50nM, 3h) and various concentrations of RAP normalized to cells treated with PBS (n=3, 10000 cells each). Statistical significance is determined by two-way ANOVA with uncorrected Fisher’s LSD multiple comparison tests. Displayed is the multiple comparison of cells treated with 500 nM RAP against cells with no treatment for each cell line. **(F)** Representative images of iPSC-derived astrocytes immunolabeled with anti-10E4, an HSPG marker. Green, 10E4; blue, Hoechst 33342; scale bar = 10 µm **(G)** Quantification of the expression of HSPGs using fluorescence intensity of anti-10E4 normalized to that of Hoechst per field of view (n=24 FOVs taken from 8 biological replicates, 3 FOVs per biological replicate) **(H)** Quantification of the expression of HSPG-related genes with RNA sequencing (n=3 for WT and n=6 for APOE3-Ch). Z-score of each HSPG gene is color coded and the size of the circle represents the mean expression. P-values were calculated using a Wald test for differential expression between groups (WT vs APOE3-Ch). Multiple testing correction was performed using the Benjamini-Hochberg method to control the false discovery rate (FDR). Genes with an adjusted p-value < 0.05 were considered statistically significant. Significantly upregulated genes were marked by red asterisks and significantly downregulated genes were marked by blue asterisks. **(I)** Representative images of iPSC-derived astrocytes immunolabeled with anti-LRP1. Green, LRP1; blue, Hoechst 33342; scale bar = 10 µm **(J)** Quantification of the expression of LRP1 using fluorescence intensity of anti-LRP1 normalized to that of Hoechst per field of view (n=12 FOVs taken from 4 biological replicates, 3 FOVs per biological replicate) All data are expressed as mean ± s.d. with individual data points shown. One-way ANOVA with Dunnett’s multiple comparison tests was performed to determine the significance unless otherwise specified. The results designated as “ns” are not significant; **p<0.01; ***p<0.001;****p<0.0001

To explore the mechanisms underlying tau uptake in astrocytes, we conducted competition assays using heparin, which inhibits tau uptake by binding to heparan sulfate proteoglycans (HSPGs) (Fig.3D). In WT astrocytes, tau uptake showed only a slight decrease in the presence of heparin, suggesting that HSPGs play a minimal role in mediating tau internalization. The two APOE3-Ch clones (1262 and 1264) exhibited variability in their response to heparin inhibition, possibly due to clonal differences in iPSC lines affecting sensitivity to HSPGs. However, both APOE3-Ch clones inhibited HSPG-mediated uptake when 100 μg/ml of heparin was co-incubated with tau oligomers, suggesting a prominent role of HSPGs in facilitating tau oligomer uptake in APOE3-Ch astrocytes.

We also assessed the role of LRP1 in tau uptake using RAP, an LRP1 antagonist (Fig.3E). RAP treatment drastically reduced tau uptake in both WT and APOE3-Ch astrocytes, illustrating the essential role of LRP1 in tau internalization. While LRP1 is critical in both astrocyte types, HSPG-mediated tau uptake appears more involved in APOE3-Ch astrocytes, potentially contributing to their increased tau internalization.

To further elucidate the mechanisms underlying differential tau uptake, we measured the expression of LRP1 and HSPGs via immunocytochemistry. APOE3-Ch astrocytes exhibited significantly higher HSPG expression, correlating with the greater reliance on HSPG-mediated tau uptake (Fig.3, F and G). Corroborating the immunocytochemistry findings, our RNA sequencing data also suggest APOE3-Ch astrocytes increase HSPG expression by significantly upregulating the expression of several HSPG-related genes including B3GAT1, EXT1, EXTL2, GPC3, HS3ST3A1, HSPG2, NDST1, NDST3, SDC2, SULF1, SULF2, TGFBR3 and XYLT1 (Fig.3H). In contrast, LRP1 expression did not differ significantly between APOE3-Ch and WT astrocytes (Fig.3, I and J). These findings suggest that APOE3-Ch astrocytes may facilitate tau uptake through both LRP1- and HSPG-mediated pathways. The increased expression of HSPGs, combined with the use of these dual uptake mechanisms, results in enhanced tau internalization in APOE3-Ch astrocytes compared to wild-type astrocytes. This differential uptake reflects an intrinsic characteristic of APOE3-Ch astrocytes, potentially contributing to their protective role against Alzheimer’s Disease.

### APOE3-Christchurch Astrocytes Promote Tau Clearance by Degradation, Not Secretion

After characterizing tau uptake, we investigated tau clearance mechanisms in APOE3-Ch astrocytes. Cells typically clear proteins via degradation or secretion pathways. Based on patient studies showing limited tau pathology in the individual carrying the APOE3-Ch variant and the familial Alzheimer’s Disease mutation PSEN1-E280A (*6*), we hypothesized that the APOE3-Ch variant enhances astrocytic tau clearance and thereby limits tau spread in the brain. We conducted tau clearance assays by incubating astrocytes with fluorescently tagged 2N4R-P301L tau oligomers, followed by media change and wash steps to remove non-internalized tau. We then measured intracellular and extracellular tau levels over time via flow cytometry (Fig.4A).

**Fig.4:**
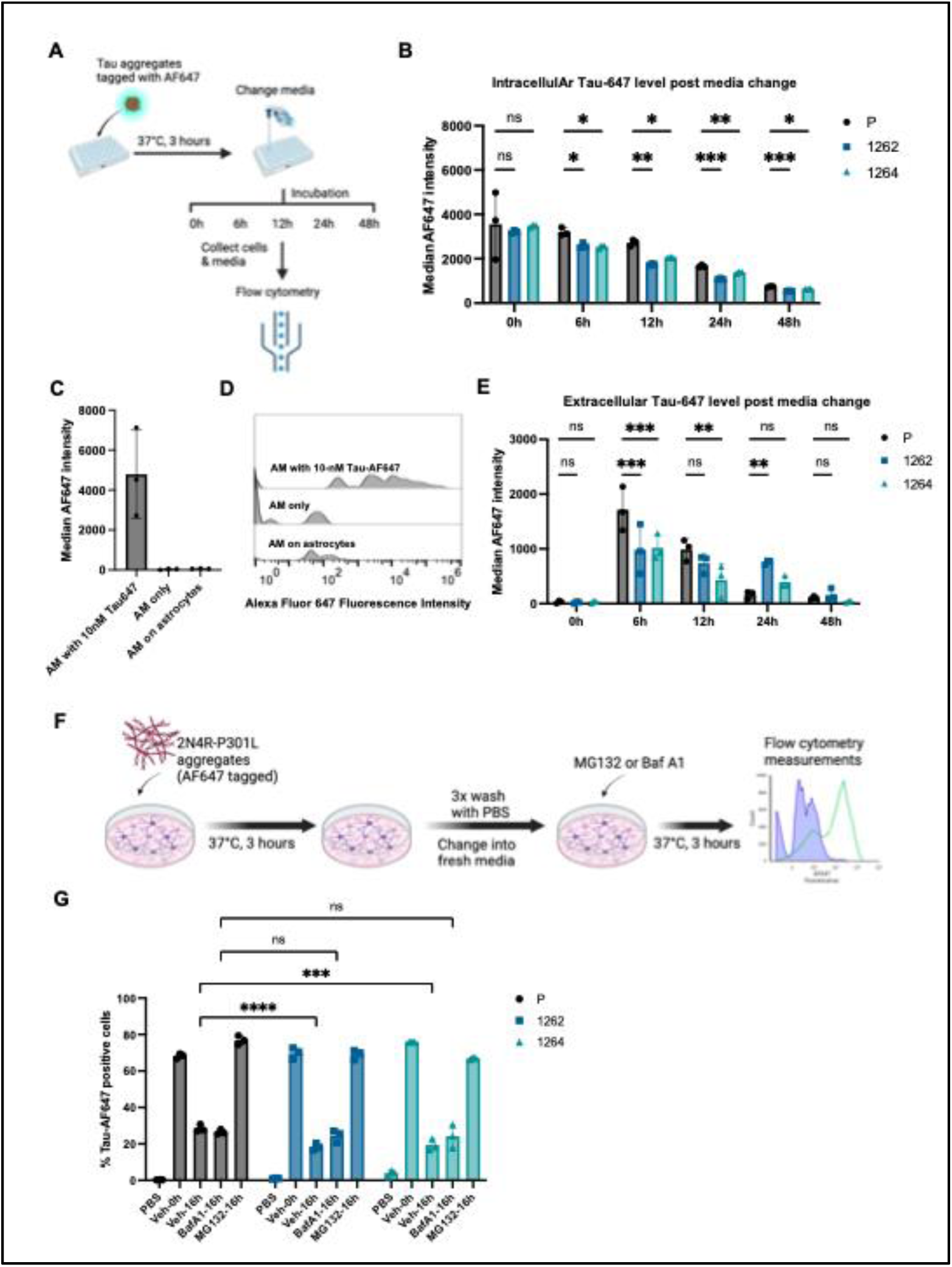
APOE3-Ch astrocytes promote more tau degradation and less secretion compared to WT astrocytes. **(A)** Schematic of the Tau clearance assay in iPSC-derived astrocytes **(B)** Measurements of the intracellular Tau-AF647 level post media change by flow cytometry at indicated time points (n=3, 10000 cells each). Statistical significance is determined by two-way RM ANOVA with Dunnett’s multiple comparison tests. **(C)** Measurement of the median AF647 intensity in the control samples by flow cytometry. The controls include a positive control, astrocyte media (AM) containing 10nM of Tau-AF647, and two negative controls, astrocyte media only and astrocyte media placed on astrocytes for 1 day (n=3) **(D)** Representative overlaid histograms demonstrating the shift in fluorescence of media containing 10 nM of Tau-AF647 compared to the negative controls. **(E)** Measurement of the extracellular Tau-AF647 level in the media of cells treated with Tau-AF647 (100nM, 3h) after media change at 0h, 6h, 12h, 24h, and 48h (n=3). Statistical significance is determined by two-way RM ANOVA with Dunnett’s multiple comparison tests. **(F)** Schematic of the Tau clearance assay with inhibitors of the protein degradation pathways, including Bafilomycin A1 and MG132. **(G)** Measurements of the intracellular Tau-AF647 level post media change with or without the treatment of MG132 (15µM, 16h), a proteosome inhibitor, and Bafilomycin A1 (0.5µM, 16h), a V-ATPase inhibitor preventing the acidification of endosomes and lysosomes, by flow cytometry in cells treated with Tau-AF647 (100nM, 3h, n=3, 10000 cells each). All data are expressed as mean ± s.d. with individual data points shown. The results designated as “ns” are not significant; *p<0.05; **p<0.01; ***p<0.001;****p<0.0001.

APOE3-Ch astrocytes demonstrated a more pronounced reduction in intracellular tau levels, persisting from 6 to 48 hours after media change, suggesting enhanced tau clearance (Fig.4B and fig.S6A). During the first 12 hours post media change, APOE3-Ch astrocytes exhibited a faster clearance rate compared to WT astrocytes, while the opposite was observed from 12 to 48 hours (fig.S6, B and C). This indicates that APOE3-Ch astrocytes activate tau clearance mechanisms more rapidly.

To elucidate the mechanisms behind this rapid clearance, we performed colocalization studies with markers of lysosomal degradation (LAMP1, LC3B, and P62) and proteasomal degradation (20S proteasome). At 8 hours post-media change, APOE3-Ch astrocytes showed increased colocalization of tau with LAMP1, LC3B, and P62, suggesting an early engagement of lysosomal degradation (fig.S4, A-D and fig.S5, A-H). By 16 hours, this trend was diminished or reversed. Colocalization with the 20S proteasome followed a different pattern; no significant differences were observed at 8 hours, but increased colocalization was detected in APOE3-Ch (1262) astrocytes at 16 hours (fig.S4, E-H). These findings indicate that APOE3-Ch astrocytes initially rely more on lysosomal degradation for tau clearance.

We also examined extracellular tau levels to determine if enhanced clearance was due to increased secretion (Fig.4C-E). APOE3-Ch astrocytes exported less tau compared to WT astrocytes (Fig.4E), indicating that the enhanced clearance is primarily mediated by degradation rather than secretion.

We further validated these findings using tau degradation inhibition assays with MG132, a proteasome inhibitor, and Bafilomycin A1 (BafA1), a lysosome inhibitor (Fig.4F). Before treatment (Veh-0h), APOE3-Ch astrocytes contained more internalized tau compared to WT astrocytes, consistent with previous findings that APOE3-Ch astrocytes can take up more tau (Fig.4G). After 16 hours of incubation with MG132, both WT and APOE3-Ch astrocytes showed a halt in tau clearance, indicating that proteasomal degradation is essential for tau clearance in astrocytes. Interestingly, treatment with Baf A1 did not completely halt tau clearance but abolished the differences in tau clearance between APOE3-Ch and WT astrocytes, resulting in comparable tau retention in both cell types. Notably, Baf A1 treatment had minimal impact on tau clearance in WT astrocytes, while it partially inhibited tau clearance in APOE3-Ch astrocytes, suggesting that lysosomal degradation contributes more specifically to the enhanced tau clearance observed in APOE3-Ch astrocytes.

Finally, we confirmed these results using total tau ELISA at representative time points (fig.S6, D and E). After 24 hours of clearance, the total tau concentration in APOE3-Ch astrocytes decreased approximately twofold, whereas it did not change significantly in WT astrocytes. Additionally, the extracellular tau concentration in APOE3-Ch astrocytes was lower than that in WT astrocytes after 24 hours, supporting our observation that APOE3-Ch astrocytes favor degradation over secretion and tau is less likely to spread from APOE3-Ch astrocytes

Overall, while proteasomal degradation is critical in both APOE3-Ch and WT astrocytes, there is no evidence to suggest that proteasomal activity differs significantly between the two astrocyte types. These results suggest that the enhanced tau clearance observed in APOE3-Ch astrocytes is partly due to their effective utilization of lysosomal pathways.

### Transcriptomic Analysis Unveils Enhanced Cell Projection and Endocytosis in APOE3Ch Astrocytes for Tau Clearance

To investigate the effects of excess tau on astrocytes and the differences between wild-type (WT) and APOE3-Ch astrocytes at both basal levels and upon tau treatment, we performed RNA sequencing on four groups: WT untreated, WT treated with tau, APOE3Ch untreated, and APOE3Ch treated with tau (Fig.5A).

**Fig.5:**
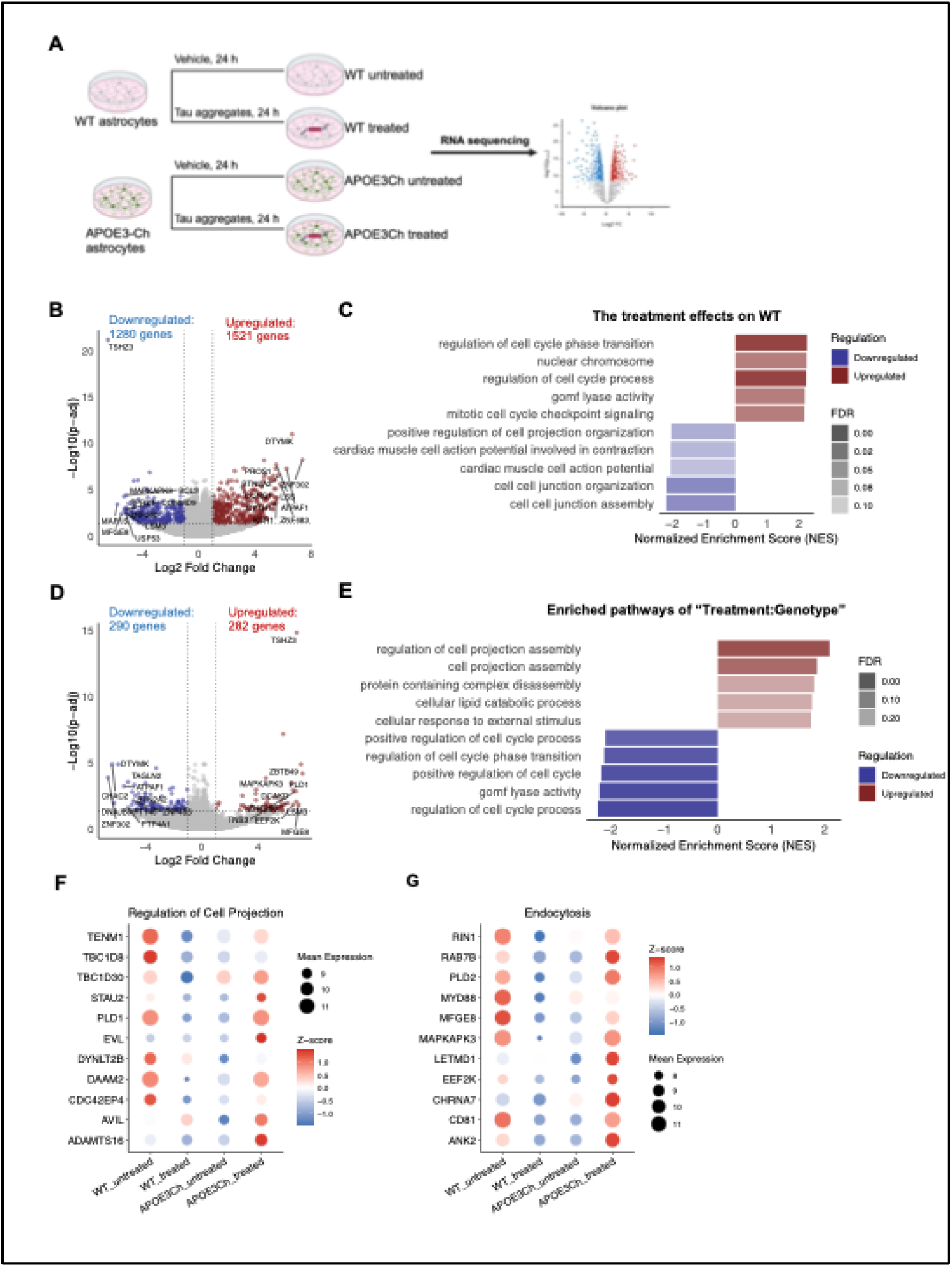
Transcriptomics studies reveal APOE3-Ch astrocytes upregulate genes related to cell projection and endocytosis upon tau treatment. **(A)** Schematics of the samples processed by RNA sequencing **(B)** Volcano plot showing differentially expressed (DE) genes in WT-treated astrocytes compared to WT-untreated astrocytes. Statistically significant (Padj < 0.05, |Log2 fold change| > 1) DE genes are labeled in red (upregulated) and blue (downregulated). The top 10 upregulated and downregulated genes ranked by their fold change are labeled with their corresponding gene symbols. **(C)** Gene set enrichment analysis (GSEA) results of the top upregulated pathways (red) and downregulated pathways (blue) ranked by their normalized enrichment score (NES) comparing WT-treated astrocytes to WT-untreated astrocytes. Pathways with a false discovery rate (FDR) < 0.1 are shown and the FDR of each pathway is represented on a color transparency scale. **(D)** Volcano plot showing differentially expressed (DE) genes in the interaction term, “Treatment:Genotype”. Statistically significant (Padj < 0.05, |Log2 fold change| > 1) DE genes are labeled in red (upregulated) and blue (downregulated). The top 10 upregulated and downregulated genes ranked by their fold change are labeled with their corresponding gene symbols. **(E)** Gene set enrichment analysis (GSEA) results of the top upregulated pathways (red) and downregulated pathways (blue) ranked by their normalized enrichment score (NES) in the interaction term, “Treatment:Genotype”. Pathways with a false discovery rate (FDR) < 0.3 are shown and the FDR of each pathway is represented on a color transparency scale. **(F)** Quantification of the mean expression and z-score of each DEG (p.adj < 0.05, abs(Log2FC) >= 1) involved in the GO Biological Process “Regulation of Cell Projection”. The size of the dot represents the mean expression, and the z-score is plotted on a color gradient. **(G)** Quantification of the mean expression and z-score of each DEG (p.adj < 0.05, abs(Log2FC) >= 1) involved in the GO Biological Process “Endocytosis”. The size of the dot represents the mean expression, and the z-score is plotted on a color gradient.

Differential gene expression analysis provided insights into how tau oligomers impacted WT and APOE3Ch astrocytes, revealing distinct responses between the two genotypes.

First, to understand the effects of tau treatment on WT astrocytes, we compared the differentially expressed (DE) genes between WT-treated and WT-untreated astrocytes (Fig.5B). Gene set enrichment analysis indicated that the top-upregulated pathways were primarily involved in the cell cycle process (Fig.5C), potentially indicating a stress response leading to increased cellular proliferation as repair mechanisms in WT astrocytes under tau stress.

Given our previous findings that APOE3Ch astrocytes process tau differently from WT astrocytes, we investigated the genotype-treatment interaction in the RNA sequencing data to compare their transcriptomic responses to tau treatment (Fig.5, D and E). The genotype-treatment interaction term measures how the effects of tau treatment in APOE3-Ch astrocytes differ from those in WT astrocytes, relative to each genotype’s no-treatment condition. This interaction analysis revealed that, upon tau treatment, APOE3-Ch astrocytes upregulate genes involved in cell projection assembly, while WT astrocytes showed downregulation in this process, along with concurrent upregulation of cell cycle-related genes. Astrocyte cell projections, or extensions, are critical for cellular communication (*26*) and for the clearance of toxic oligomers such as tau (*27*).

Close examination (Fig.5F) revealed that several key genes involved in regulating cell assembly of cell projections, including TENM1, TBC1D8, TBC1D30, STAU2, PLD1, DYNLT2B, DAAM2, CDC42EP4, and ADAMTS16, were downregulated in tau-treated WT astrocytes. In contrast, these genes were upregulated in APOE3-Ch astrocytes following tau treatment, suggesting that APOE3-Ch astrocytes enhance their ability to form projections, possibly to aid in the clearance of tau oligomers.

In parallel, genes associated with the endocytosis pathway, such as RIN1, RAB7B, PLD2, MYD88, MFGE8, MAPKAPK3, LETMD1, EEF2K, CHRNA7, CD81, and ANK2, also followed a similar pattern (Fig.5G). These genes are involved in all aspects of endocytosis. For instance, RIN1 and PLD2 have been implicated in activating receptor-mediated endocytosis (*28*, *29*), RAB7B in late endosomes (*30*, *31*), and CD81 in exosomes and multivesicular bodies (MVBs) (*32*). APOE3-Ch astrocytes upregulated these genes upon tau treatment, suggesting an enhancement of endocytosis and subsequent lysosomal degradation pathways. This aligns with our prior findings that APOE3-Ch astrocytes show enhanced tau internalization and degradation through the lysosomal pathway.

In summary, our transcriptomic analysis reveals a complex and coordinated defense mechanism in APOE3-Ch astrocytes in response to tau pathology. This involves upregulation of cell projections and endocytosis pathways, facilitating enhanced uptake and degradation of tau, which may confer protective effects against tau toxicity.

## Discussion

In this study, we characterized the phenotypic and functional differences between wild-type (WT) and APOE3-Christchurch (APOE3-Ch) astrocytes, uncovering several key differences in lipid metabolism, tau uptake, and clearance mechanisms. We found that APOE3-Ch astrocytes exhibit lipid dyshomeostasis, characterized by decreased lipid buffering capacity and impaired APOE secretion. Lipidomic analysis revealed significantly reduced ceramide levels in APOE3-Ch astrocytes, which is likely driven by the upregulation of ceramide catabolic pathways. Our analysis of tau uptake demonstrated that APOE3-Ch astrocytes internalize tau oligomers more efficiently than WT astrocytes, a process mediated by both HSPG and LRP1 pathways. Following tau internalization, APOE3-Ch astrocytes degraded tau through both lysosomal and proteasomal pathways, while WT astrocytes primarily relied on proteasomal degradation and secreted more tau than APOE3-Ch astrocytes. This decrease in secreted tau in APOE3-Ch astrocytes may contribute to the limited tau spread in the human case expressing the Christchurch variant (*10*). Finally, transcriptomic analysis showed upregulation of cell projection and endocytic pathways in APOE3-Ch astrocytes, the latter potentially enhancing their ability to internalize and degrade tau oligomers. These transcriptional changes align with our cellular assay findings and suggest that APOE3-Ch astrocytes have a heightened capacity for clearing tau.

APOE3-Ch is a rare genetic variant of APOE and has been associated with hyperlipoproteinemia type III (HLPP3) in patients (*18*). APOE3-Ch mice have exhibited peripheral dyslipidemia, including elevated APOE and APOB levels, as well as increased association of APOE with very-low-density lipoproteins (VLDL) and low-density lipoproteins (LDL) (*8*). However, the effects of APOE3-Ch on central nervous system (CNS) lipid metabolism remain unclear. Our findings suggest that similar to astrocytes expressing APOE4—an APOE variant known to increase risk for Alzheimer’s disease (AD) and cardiovascular diseases—APOE3-Ch astrocytes show reduced capacity to buffer lipids (*33*). However, unlike APOE4 astrocytes, APOE3-Ch astrocytes do not accumulate unsaturated triacylglycerols (TAGs), which are prone to lipid peroxidation (*34*). The accumulation of unsaturated TAGs in APOE4 astrocytes may promote oxidative stress and pro-inflammatory profiles, contributing to neuroinflammation and neurotoxicity (*34*), processes that are not observed in APOE3-Ch astrocytes.

Additionally, our data show that APOE3-Ch astrocytes have reduced levels of ceramides both intracellularly and in the media. Ceramides are bioactive lipids that play a significant role in several pathological processes associated with AD, including amyloid-beta plaque formation, mitochondrial dysfunction, cellular senescence, and autophagy dysfunction (*21*). Ceramides are also known to initiate some of the pathological changes that drive AD progression (*21*, *22*). The lower ceramide levels observed in APOE3-Ch astrocytes may explain, in part, why this variant is protective against AD, despite the observed lipid dyshomeostasis. Future studies should investigate how specific ceramide species influence astrocyte function.

The role of astrocytic APOE in tau-mediated neurodegeneration has been highlighted by recent work showing that selective removal of astrocytic APOE4 strongly protects against tau-induced synaptic loss and reduces microglial-mediated synaptic phagocytosis (*35*). This underscores the isoform-specific effects of APOE on tau pathology and neurodegeneration. While APOE4 impairs astrocytic function and exacerbates tau toxicity, our findings suggest that APOE3-Ch astrocytes confer protection by enhancing tau clearance via HSPG- and LRP1-mediated pathways. Previous studies have shown that astrocytes can internalize tau monomers (*13*), truncated pre-formed fibrils (PFFs) (*12*), and tau oligomers (*11*, *36*) through HSPG-independent mechanisms (*25*). Our data suggest that, like neurons, which rely heavily on LRP1-mediated and HSPG-mediated uptake (*37–39*), APOE3-Ch astrocytes, utilize both HSPG- and LRP1-mediated pathways for tau oligomer internalization, while WT astrocytes primarily rely on LRP1-mediated tau uptake. The dual mechanisms of LRP1- and HSPG-mediated uptake in APOE3-Ch astrocytes are reminiscent of their synergistic roles in Aβ uptake observed in primary neurons (*40*). Research indicates that LRP1 can form complexes with HSPGs, allowing ligands such as APOE to either bind directly to the LRP1-HSPG complex or initially interact with HSPGs before being transferred to LRP1 for endocytosis (*41*). However, our study does not address whether HSPGs and LRP1 cooperate directly in tau uptake in APOE3-Ch astrocytes or elucidate the involvement of specific endocytosis pathways, such as macropinocytosis or receptor-mediated endocytosis. The dual engagement of these two mechanisms represents a novel protective feature of the APOE3-Ch variant, allowing for more efficient tau uptake and clearance from the extracellular space.

Following tau internalization, our data show that APOE3-Ch astrocytes degrade tau primarily through lysosomal and proteasomal pathways, with lysosomal degradation playing a particularly vital role. In contrast, WT astrocytes rely more heavily on proteasomal degradation and secrete a greater proportion of tau. Lysosomal degradation in APOE3-Ch astrocytes occurs more rapidly, leading to enhanced tau clearance and preventing the accumulation of tau oligomers. The transcriptomic upregulation of endocytic and cell projection pathways in APOE3-Ch astrocytes aligns with these findings, as these pathways likely enhance the cells’ ability to internalize and degrade tau. By scavenging tau oligomers more effectively, APOE3-Ch astrocytes may prevent the spread of tau pathology and mitigate neurotoxicity.

Our findings contribute to the growing understanding of APOE3-Ch as a protective variant against AD. Unlike the APOE4 variant, which has been associated with both impaired lipid metabolism and tau-induced neurotoxicity, APOE3-Ch astrocytes seem to offer protection through two mechanisms: enhanced tau clearance and reduced harmful lipid levels. The enhanced tau uptake and degradation observed in APOE3-Ch astrocytes may help to prevent tau accumulation and spreading, a hallmark of AD progression. The reduced ceramide and GLA levels provide further protection by mitigating the bioactive lipid-mediated signaling pathways that exacerbate AD pathology.

Although our study highlights these protective mechanisms, the observed lipid dysregulation in APOE3-Ch astrocytes suggests that this variant’s effects are complex. While the variant helps clear tau and reduce neurotoxic lipids like ceramides, it may also impair normal lipid metabolism, potentially leading to long-term vulnerabilities in astrocyte-neuron metabolic coupling. Future studies should aim to better understand how the lipid dysregulation we observed may influence astrocyte function over time and whether it poses risks for other neurodegenerative conditions.

A limitation of our study is the use of iPSC-derived astrocytes, which may not fully replicate the mature astrocyte phenotype observed in vivo. For example, the low expression of AQP4 in our astrocytes may suggest incomplete maturation, which could affect the generalizability of our findings. Future research should focus on validating these results in more mature astrocyte models or in vivo systems to ensure that the observed phenotypes hold true in fully developed astrocytes.

Additionally, further investigation is needed to understand the mechanisms underlying the reduced exported APOE observed in APOE3-Ch astrocytes. Given the central role of APOE in lipid transport and metabolism, understanding how this reduction impacts lipid handling and its broader implications for brain health could provide new insights into the role of APOE variants in neurodegenerative diseases. Finally, future studies should explore the effects of APOE3-Ch astrocytes on neuronal function, particularly in co-culture or animal models, to better understand how these cells influence tauopathy and other neurodegenerative processes.

## Materials and Methods

### Experimental Design

This study investigates the protective mechanisms conferred by the APOE3-Christchurch (APOE3-Ch) variant in the context of Alzheimer’s disease (AD), focusing on astrocytic lipid metabolism and tau processing. To achieve this, human-induced pluripotent stem cell (hiPSC)-derived astrocytes from a wild-type (WT) line and two lines homozygous for the APOE3-Ch variant were utilized. The objectives included characterizing differences in lipid metabolism, elucidating the mechanisms underlying tau uptake and clearance, and identifying transcriptomic changes linked to protective pathways.

Lipid metabolism was analyzed using a combination of lipidomic profiling, lipid droplet induction, and APOE secretion assays. Lipid droplet formation was induced using oleic acid, and the resulting droplets were quantified using confocal microscopy and computational image analysis. Lipidomic profiling was performed on cell and media samples to assess intracellular and secreted lipid species, and APOE secretion was quantified using enzyme-linked immunosorbent assay (ELISA).

Tau processing and degradation mechanisms were studied using purified, fluorescently labeled tau oligomers to assess uptake and degradation pathways. Competition assays with heparin and RAP determined the roles of HSPGs and LRP1 in tau uptake.

Degradation pathways were further investigated using lysosomal and proteasomal inhibitors, followed by flow cytometric quantification of tau clearance.

Transcriptomic profiling was conducted to identify the molecular pathways associated with APOE3-Ch astrocytes. RNA sequencing was performed on astrocytes under basal conditions and following tau treatment, enabling differential expression analysis and pathway enrichment analysis.

All experiments were conducted in biological triplicates or quadruplicates to ensure robustness and reproducibility.

### Ethical Statement

Our research complies with all relevant ethical regulations sat the University of California Santa Barbara and study procedures were approved by the Institutional Review Board (#767).

### hiPSC Culture

Human induced pluripotent stem cell (hiPSC) lines, including a WT parental line (KOLF2.1J, Cat. No. JIPSC1000) and two lines homozygous for the APOE3-Ch mutation (denoted as 1262 and 1264, Cat. No. JIPSC1262 and JIPSC1264), were obtained from The Jackson Laboratory. Cells were cultured in Matrigel-coated (Corning, Cat. No. 354277), tissue-culture-treated plates using mTeSR Plus medium (Stemcell Technologies, Cat. No. 100-0276). The medium was replaced every other day until the cells reached approximately 70% confluence. Cells were passaged at a 1:20 ratio using ReLeSR (Stemcell Technologies, Cat. No. 100-0483), following the manufacturer’s protocol. Cells were cryopreserved in mTeSR Plus medium with 10% DMSO (Sigma-Aldrich, Cat. No. D8418).

### Differentiation of iPSCs into Astrocytes

iPSC-derived astrocytes were differentiated from both WT and APOE3-Ch iPSCs using a modified version of previously published protocols (Pantazis et al., 2022; TCW et al., 2017) were dissociated using Accutase (Stemcell Technologies, Cat. No. 07922) and seeded into 24-well AggreWell 800 plates (Stemcell Technologies, Cat. No. 34815) at a density of 2 million cells per well in neural induction medium, which consists of DMEM/F-12 (Gibco, Cat. No. 11320033), 1× B-27 supplement minus vitamin A (50X, Gibco, Cat. No. 12587010), 1× N-2 supplement (100X, Gibco, Cat. No. 17502048) and 10 nM Y-27632 Dihydrochloride or ROCK inhibitor (Stemcell Technologies, Cat. No. 72304). This setup was used to generate embryoid bodies (EBs). On Day 2, the medium was replaced with the neural induction medium containing 0.1 µM LDN193189 (Tocris, Cat. No. 6053) and 10 µM SB431542 (Tocris, Cat. No. 1614), while the ROCK inhibitor was removed. Half media changes were performed every other day.

By Day 7, EBs exhibited clear and circular borders. The EBs were collected and plated onto Matrigel-coated plates. Adherent cultures were maintained with the same media and fed every other day for an additional week. On Day 14, neural rosettes were isolated using STEMdiff Neural Rosette Selection Reagent (Stemcell Technologies, Cat. No. 05832) and replated on Matrigel-coated plates in neural progenitor cell (NPC) medium, which consists of DMEM/F12, 1× B27 minus vitamin A, 1× N2 supplement, and 20 ng/ml FGF-2 (Thermo Fisher Scientific, Cat. No. PHG0367) in 1% BSA (Sigma-Aldrich, Cat. No. A9418). Full media changes were conducted every other day until confluency was reached.

Around Day 21, NPCs were harvested and stained with PE-conjugated anti-CD133/1 antibody (Miltenyi Biotec, Cat. No. 130-113-670) and PerCP-Cy5.5-conjugated anti-CD271 antibody (BD Biosciences, Cat. No. 560834). CD133^+^/CD271^−^ populations were sorted using a SONY MA900 cell sorter, with mouse IgG1κ (BD Biosciences, Cat. No. 550795) as an isotype control.

CD133^+^/CD271^−^ NPCs were expanded in NPC media and cryopreserved in NPC media supplemented with 10% DMSO. Additionally, cells were plated on Matrigel-coated plates in Astrocyte Media (ScienCell Research Laboratories, Cat. No. 1801) for astrocyte differentiation. The cells were cultured in Astrocyte Media for two weeks before conducting cellular assays. Astrocytes were cryopreserved in Astrocyte Media with 10% DMSO for later use.

### RNA Extraction and Sequencing

Differentiated astrocytes were plated in 60-mm dishes and cultured until they reached full confluency. Three biological replicates were prepared for each cell line, including three replicates for WT astrocytes and three replicates each for the 1262 and 1264 cell lines. For RNA extraction, cells were lysed with TRIzol reagent (Invitrogen, Cat. No. 15596026) according to the manufacturer’s instructions. Cold chloroform (Sigma-Aldrich, Cat. No. 366927) was added to the TRIzol cell suspension at a ratio of 1:5. The mixture was vortexed for 15 seconds and then incubated at room temperature for 3 minutes to allow for phase separation. Following incubation, the sample was centrifuged at 12,000 × g for 15 minutes at 4°C. The transparent aqueous (upper) phase was carefully transferred to a new tube, ensuring no disruption of the interphase layer. An equal volume of isopropanol (Sigma-Aldrich, Cat. No. 34863) was added to the aqueous phase, and the tube was gently inverted five times to mix. The mixture was then incubated at room temperature for 10 minutes, followed by centrifugation at 12,000 × g for 10 minutes at 4°C. The supernatant was discarded. The RNA pellet was washed by adding an equal volume of cold ethanol (Sigma-Aldrich, Cat. No. 459828), followed by brief vortexing. The sample was then centrifuged at 7,500 × g for 5 minutes at 4°C. The supernatant was thoroughly removed without disturbing the RNA pellet. The RNA pellet was resuspended in UltraPure DNase/RNase-free distilled water (Invitrogen, Cat. No. 10977015) and incubated at 55°C for 10 minutes with gentle vortexing every 5 minutes to ensure complete dissolution.

Library preparation for RNA sequencing using NEBNext Ultra II Directional RNA Library Prep Kit (New England Biolabs, Cat. No. E7760S) with the NEBNext Poly(A) mRNA Magnetic Isolation Module (New England Biolabs, Cat. No. E7490S) and NEBNext Multiplex Oligos for Illumina (New England Biolabs, Cat. No. E7335S) and Next-Generation Sequencing were performed at the UCSB Biological Nanostructures Laboratory. Sequencing was done on an Illumina NextSeq 500 using a 75-cycle high-output kit.

### Transcriptomic Analysis

RNA sequencing fastq files were quality-checked using FastQC and trimmed with Trimmomatic. The trimmed fastq files were then aligned to the GRCh38 human genome assembly with HISAT2. FeatureCounts was used to generate a count matrix by counting gene features. Differential expression analysis was conducted with the DESeq2 pipeline, which combines the two APOE3-Ch lines to represent a single genotype and compares phenotypic differences between genotypes. Gene counts were normalized using the DESeq2 rlog function for regularized log transformation.

### Lipid Droplet Induction and Quantification

To induce lipid droplet production and assess lipid buffering capacity in astrocytes, oleic acid (Sigma-Aldrich, Cat. No. O1008) was dissolved in a fatty acid-free BSA solution in 0.1M Tris, pH 8.0 (Invitrogen, Cat. No. AM9855G), following a previously published protocol (Brasaemle and Wolins, 2016). Differentiated astrocytes were plated at a density of 20,000 cells per well on Matrigel-coated 96-well glass-bottom plates in Astrocyte Media. Cells were incubated with 20 µM oleic acid in BSA at 37°C for 18 hours. After incubation, cells were fixed with 4% paraformaldehyde and stained with LipidTOX Deep Red (Invitrogen, Cat. No. H34477, 1:1000) and Hoechst 33342 (same as above, 1:10000) at room temperature for 30 minutes. Following staining, cells were washed twice with PBS and stored in PBS until imaging. Imaging was conducted with a Leica SP8 resonant scanning confocal microscope at the UCSB NRI-MCDB Microscopy Facility, using a 60X objective.

Lipid droplet quantification was conducted using a puncta identification algorithm. Puncta centers were detected using the blob_log function from the scikit-image library (version 0.24.0), which implements the Laplacian of Gaussian method for blob detection. The minimum and maximum size parameters for puncta were determined based on manual measurements performed in ImageJ. To ensure detection quality, each identified punctum was enclosed within a bounding box of 2σ width (where σ values for each punctum are provided by blob_log), and puncta with a signal-to-noise ratio below 1% were excluded to remove low-quality detections. The shape of each punctum was further refined using a flood-fill function applied to the inverted intensity values within the bounding box. This approach enabled the extraction of detailed shape characteristics, including area, eccentricity, and peak intensity.

### APOE ELISA

120,000 differentiated astrocytes were plated in Matrigel-coated 24-well plates prior to collection. On the collection day, media were harvested and centrifuged at 1500g for 5 minutes to remove cell debris, with the supernatants retained as media samples. Cell pellets were then lysed in RIPA buffer (Thermo Fisher Scientific, Cat. No. 89900) supplemented with EDTA-free cOmpleteTM protease inhibitor cocktail (Roche, Cat. No. 11836170001) and 1 mM PMSF (0.1 M, Boston BioProducts, Cat. No. BP-481). The total protein concentration of both cell pellet samples and media samples was determined using the PierceTM BCA protein assay kit (Thermo Fisher Scientific, Cat. No. 23225). APOE quantification was performed using the Human APOE ELISA kit (Invitrogen, Cat. No. EHAPOE), following the manufacturer’s protocol. The normalized APOE concentration for both cell pellet and media samples was calculated by normalizing the APOE concentration measured by ELISA to the total protein concentration of the corresponding cell pellet sample.

### Cell Viability Assay

Differentiated astrocytes were plated in a Matrigel-coated 96-well plate at a density of 20,000 cells per well. Control wells containing medium without cells were included to determine background luminescence. The next day, cells were treated with 20 µM oleic acid in BSA for 18 hours. Cell viability was assessed using the CellTiter-Glo Cell Viability Assay (Promega, Cat. No. G7570) according to the manufacturer’s instructions. Briefly, the plate and its contents were equilibrated to room temperature, and CellTiter-Glo Reagent was added directly to wells containing medium. The plate was then mixed on a shaker for 2 minutes to induce cell lysis and incubated at room temperature to stabilize the luminescent signal. Luminescence was measured using a Tecan Spark 10M Multimode Plate Reader.

### Lipidomics

Differentiated astrocytes were plated in 10-cm dishes and cultured to full confluency. Four biological replicates were prepared for each cell line, including four replicates for WT astrocytes and four replicates each for the 1262 and 1264 cell lines. Media were collected and centrifuged at 1500g for 5 minutes. Cell pellets were dissociated with Accutase by incubation at 37°C for 10 minutes, followed by centrifugation at 2000g for 5 minutes at 4°C. Both media samples and cell pellets were frozen at -80°C and lyophilized. The lyophilized samples were processed at the University of Texas Medical Branch Mass Spectrometry Facility for targeted lipidomics of neutral lipids, polar lipids, and fatty acids.

For sample processing, isotopically labeled lipid standards (UltimateSPLASH ONE) and free fatty acids (FFAs) were added to each sample prior to extraction. Samples were extracted with hexane (3:2, v/v), centrifuged, and the supernatant was evaporated under nitrogen. Dried samples were resuspended in 100 µL dichloromethane/methanol (1:1, v/v). Protein concentration of the remaining cell pellets was determined using a BCA assay. For lipidomics, 30 µL of each sample extract was transferred into a glass autosampler vial for neutral lipid analysis, and 50 µL was dried, resuspended in 50 µL of 80% methanol, and derivatized with 3-NPH.

LC-MS/MS analysis was performed using an Acquity H-class HPLC System (Waters) coupled to a QTRAP 6500 mass spectrometer (SCIEX). Neutral lipids were separated by reverse-phase chromatography on an Accucore C30 column (Thermo Scientific). For triacylglycerols (TG) and determination of their fatty acid (FA) compositions, the pooled QC was injected multiple times with different survey methods to cover TG FA carbon numbers from 44 to 60. FAs analyzed included 14:0, 15:0, 16:0, 16:1, 17:0, 18:0, 18:1, 18:2, 18:3, 19:0, 20:0, 20:1, 20:2, 20:3, 20:4, 20:5, 22:4, 22:5, 22:6, 24:0, and 24:1. A Python script (v3.11.5) generated all possible TG combinations, providing precursor and product ion masses based on FA fragmentation. Overlapping TG-FA peaks were curated into a final, sample-specific method to determine the complete FA composition for each TG molecular species in the lipid extracts.

### Lipidomic Analysis

LCMS peak areas were integrated using MultiQuant (v3.0.3, SCIEX). FFA peak areas were then exported and normalized to the peak area its respective [13C6]-3NPH internal standard and protein concentration of each sample (pellet samples only) using a custom R script (R v4.3.3, RStudio v2023.12.1) to give relative values of “Normalized Peak Area / mg protein”

Prior to data analysis, a data quality check analysis was performed using a custom R Markdown document to assess the response from internal standards within samples and the pooled QC, injection order effects, retention time variability, and variability of biological lipids.

To prepare the data for statistical testing, samples were organized by genotype and replicate. A design matrix was created to compare the APOE3-Ch and WT astrocyte genotypes, and columns with zero variance were removed to ensure robust statistical analysis. Differential expression analysis was conducted using the limma package. A linear model was fitted to the data, followed by the construction of a contrast matrix to compare the APOE3-Ch samples to WT. Empirical Bayes moderation was applied to the contrast-adjusted model, and results were extracted, adjusting p-values using the Holm method. A threshold of |logFC| > 0.1 and p < 0.05 was applied to identify statistically significant lipid species.

### Tau Purification

Full-length tau protein with the P301L point mutation (2N4R-P301L) was purified following a previously published protocol(*37*). Briefly, 2N4R-P301L tau encoded in the pRK172 plasmid was expressed in E. coli BL21 (DE3). Cell pellets were collected and resuspended in a lysis buffer containing 50 mM MES (pH 6.5), 5 mM DTT, 1 mM PMSF, and 1 mM EGTA, supplemented with cOmplete protease inhibitor tablets. The cell lysate was sonicated, boiled for 10 minutes, and centrifuged at 50,000 × g for 30 minutes at 4°C. The supernatant was precipitated with 20% (w/v) ammonium sulfate and centrifuged at 20,000 × g for 30 minutes at 4°C. The resulting pellet was resuspended in 4 mL of Buffer A (50 mM MES, pH 6.5, 50 mM NaCl, 2 mM DTT, 1 mM PMSF, 1 mM EGTA) and dialyzed overnight against the same buffer. The protein was then loaded onto a HiTrap SP HP column (Cytiva, Cat. No. 17115201) and eluted using a linear NaCl gradient with Buffer B (Buffer A + 1 M NaCl). Fractions containing 2N4R-P301L tau were pooled, concentrated, and dialyzed overnight into PBS (pH 7.4). Protein concentration was determined using a BCA assay.

### Tau Labeling

2N4R-P301L tau were labeled with Alexa FluorTM 647 NHS ester (Invitrogen, Cat. No. A20006) according to the manufacturer’s instructions. After labeling, 100 mM glycine was added to quench the reaction, and unreacted dye was removed using Zeba desalting columns (Thermo Scientific, Cat. No. 89882). Average label incorporation was around 1.5 moles/mole of protein, as determined by measuring fluorescence and protein concentration (A_max_ x MW of protein/[protein] x ε_dye_).

### Tau Fibrillization

To prepare tau oligomers, monomeric 2N4R-P301L tau protein was aggregated following a previously published protocol(*37*). Briefly, 10 µM tau protein in PBS with 1 mM DTT (pH 7.4) was combined with 0.05 mg/mL heparin and incubated at 37°C with shaking for 4 hours. The resulting tau oligomers were buffer exchanged into PBS using Amicon Ultra Centrifugal Filter (Milipore, Cat. No. UFC500308) to remove excess DTT and heparin.

### Tau Uptake Assay

Astrocytes were seeded at a density of 20,000 cells per well in a 24-well plate pre-coated with Matrigel and maintained in Astrocyte Media. After 24 hours, the media were replaced, and the cells were treated with 50 nM of 2N4R-P301L tau oligomers conjugated to Alexa Fluor 647 (AF647) for 3 hours at 37°C. Following treatment, the cells were washed twice with PBS to remove any non-internalized tau and then dissociated using Accutase for flow cytometry analysis.

Dissociated cells were analyzed using a SONY MA900 cell sorter, and median AF647 fluorescence intensity was measured to quantify tau uptake. The median fluorescence intensity of each cell line treated with tau was normalized to its respective control condition (PBS treatment).

### Heparin Competition Assay

For heparin competition, astrocytes were pre-treated with media containing heparin at 0, 1, 50 and 100 μg/ml for 16 hours prior to tau treatment. Following this incubation, cells were washed twice with PBS, and fresh media containing 50 nM tau oligomers was added. The subsequent steps for the tau uptake assay were carried out as described above.

### RAP Purification

RAP was cloned into the pMCSG7 vector and purified using nickel-nitrilotriacetic acid (Ni-NTA) agarose (Qiagen, Cat. No. 30210). The RAP construct was transformed into Escherichia coli DH5α chemically competent cells (Thermo Fisher Scientific, Cat. No. 18265017). Cells were grown to an optical density (OD600) of 0.6, at which point protein expression was induced with isopropyl β-D-1-thiogalactopyranoside (IPTG).

For purification, Ni-NTA agarose was prepared with HIS binding buffer (50 mM Tris-HCl, pH 8.0, 1 M NaCl, 10 mM imidazole). Non-specific proteins were removed using HIS wash buffer (50 mM Tris-HCl, pH 8.0, 300 mM NaCl, 30 mM imidazole). RAP protein was eluted with HIS elution buffer containing 50 mM Tris-HCl, pH 8.0, 300 mM NaCl, and 300 mM imidazole. The protein concentration was quantified using the Pierce BCA protein assay (Thermo Scientific, Cat. No. 23228).

### RAP Competition Assay

For RAP competition, astrocytes were treated with recombinant RAP at 0, 50, 100 and 500 nM concurrently with tau addition. The protocol for tau uptake quantification was then performed as outlined in tau uptake assay.

### TEM

To prepare samples for TEM analysis, 5 μL of tau aggregate solution was applied to a glow-discharged copper grid (Electron Microscopy Sciences, Cat. No. FCF-200-Cu) and allowed to adsorb for 20 seconds. Excess liquid was gently removed by blotting with filter paper. The grids were then stained with 5 μL of 1.5% (w/v) uranyl acetate solution (Electron Microscopy Sciences, Cat. No. 22400-2), followed by immediate blotting to remove excess stain. A second application of 5 μL uranyl acetate was applied for 60 seconds, after which the grid was blotted dry.

The prepared grids were imaged using a Thermo Scientific Talos G2 200X TEM/STEM microscope operating at 200 kV under room temperature conditions. Images were captured using a Ceta II CMOS 4k x 4k camera at specified magnifications to visualize the tau oligomers.

### Tau Clearance Assay

Astrocytes were seeded at a density of 120,000 cells per well in a 24-well plate pre-coated with Matrigel and maintained in Astrocyte Media. After 24 hours, the media were replaced, and the cells were treated with 100 nM 2N4R-P301L tau oligomers conjugated to Alexa Fluor 647 (AF647) for 3 hours at 37°C. Following treatment, the cells were washed twice with PBS to remove any non-internalized tau. Fresh media were added to the cells, which were then incubated for 0, 6, 12, 24, and 48 hours. At each time point, cells were dissociated using Accutase, fixed with 4% paraformaldehyde at room temperature for 15 minutes, and stored in PBS at 4°C until flow cytometry analysis. Media samples were also collected at each time point, centrifuged at 1500 × g for 5 minutes to remove cellular debris, and the supernatant was stored at 4°C until analysis.

Flow cytometry was performed using a SONY MA900 cell sorter. For cell samples, gating was applied to exclude debris and isolate singlet populations, and the median AF647 fluorescence intensity of the singlet cells was measured to quantify intracellular tau levels at each time point. For media samples, controls, each with three replicates, were included: (1) a positive control containing astrocyte media with 10 nM 2N4R-P301L AF647, (2) a negative control containing only astrocyte media, and (3) astrocyte media collected from untreated astrocytes. The median AF647 fluorescence intensity for media samples was recorded over a 1-minute acquisition period.

### Tau ELISA

Astrocytes were seeded at a density of 120,000 cells per well in a 24-well plate pre-coated with Matrigel and maintained in Astrocyte Media. After 24 hours, the media were replaced, and the cells were treated with 25 nM 2N4R-P301L tau oligomers conjugated to Alexa Fluor 647 (AF647) for 3 hours at 37°C. Following treatment, the cells were washed twice with PBS to remove any non-internalized tau. Fresh media were added to the cells, which were then incubated for 0 and 24 hours. At each time point, cell pellets were lysed in RIPA buffer supplemented with EDTA-free cOmpleteTM protease inhibitor cocktail and 1 mM PMSF. Media were harvested and centrifuged at 1500g for 5 minutes to remove cell debris, with the supernatants retained as media samples. The total protein concentration of both cell pellet samples and media samples was determined using the PierceTM BCA protein assay kit. Total tau quantification was performed using the human total tau ELISA kit (Invitrogen, Cat. No. KHB0041), following the manufacturer’s protocol. The normalized tau concentration for both cell pellet and media samples was calculated by normalizing the tau concentration measured by ELISA to the total protein concentration of the corresponding cell pellet sample.

### Tau Colocalization Study

Astrocytes were seeded at a density of 20,000 cells per well in a 96-well glass bottom plate pre-coated with Matrigel and maintained in Astrocyte Media. After 24 hours, the media were replaced, and the cells were treated with 50 nM 2N4R-P301L tau oligomers conjugated to Alexa Fluor 647 (AF647) for 3 hours at 37°C. Following treatment, the cells were washed three times with PBS to remove any non-internalized tau. Fresh media were added to the cells and were then incubated for 8 and 16 hours, respectively. At each time point, cells were fixed for immunocytochemistry probing for tau colocalization with P62, LAMP1, LC3B and 20S proteasome.

### Immunocytochemistry

Differentiated astrocytes (> day 14) were plated in Astrocyte Media on Matrigel-coated 96-well glass-bottom plates (Cellvis, Cat. No. P96-0-N). Cells were fixed with 4% paraformaldehyde (16% solution, Electron Microscopy Sciences, Cat. No. 15700) at room temperature for 15 minutes, followed by three washes with ice-cold PBS (Gibco, Cat. No. 10010049). Cells were then permeabilized with 0.1% Triton™ X-100 (Sigma-Aldrich, Cat. No. X100-5ML) at room temperature for 10 minutes and blocked with 10% normal goat serum (Thermo Fisher Scientific, Cat. No. 50062Z) at room temperature for 1 hour.

For immunostaining, primary antibodies including chicken anti-GFAP (Abcam, Cat. No. ab4674, 1:500, diluted in 5% normal goat serum) and mouse anti-S100B (Invitrogen, Cat. No. MA1-25005, 1:500, diluted in 5% normal goat serum) were applied and incubated with cells overnight at 4°C on a shaker. After incubation, cells were washed with PBST (PBS + 0.1% Tween® 20) three times for 5 minutes each. Secondary antibodies, goat anti-mouse IgG1 Alexa Fluor™ 488 (Invitrogen, Cat. No. A-21121, 1:1000, diluted in 5% normal goat serum) and goat anti-chicken IgY Alexa Fluor™ 647 (Invitrogen, Cat. No. A-21449, 1:1000, diluted in 5% normal goat serum), along with Hoechst 33342 staining (Thermo Fisher Scientific, Cat. No. 62249, 1:10000, diluted in PBS), were added to the cells and incubated at room temperature in the dark on a shaker. Following incubation, cells were washed with PBST three times for 5 minutes each and stored in PBS prior to imaging.

For immunostaining of P62, LAMP1, LC3B, and 20S proteasome, the same procedure was followed. Primary antibodies were diluted in 5% normal goat serum at specific dilution factors. Mouse anti-SQSTM1/P62 (Abcam, Cat. No. ab109012) was used at a 1:500 dilution, rabbit anti-LAMP1 (Cell Signaling, Cat. No. 9091S) at 1:200, rabbit anti-LC3B (Abcam, Cat. No. ab192890) at 1:500, and rabbit anti-20S proteasome core subunits (Enzo, Cat. No. BML-PW8155-0100) at 1:500. Secondary antibody goat anti-mouse IgG Alexa Fluor 568 (Invitrogen, Cat. No. A-11004) was applied at a 1:2000 dilution in 5% normal goat serum, along with Hoechst stain.

For HSPGs and LRP1, the same procedure was followed. To probe for HSPGs, primary antibody mouse anti-10E4 (Amsbio, Cat. No. 370255-S) was applied at a 1:100 dilution in 5% normal goat serum. Secondary antibody goat anti-mouse IgG Alexa Fluor 568 (Invitrogen, Cat. No. A-11004) was applied at a 1:2000 dilution in 5% normal goat serum, along with Hoechst stain. For probing LRP1, primary antibody rabbit anti-LRP1 (Abcam, Cat. No. ab92544) was applied at a 1:1000 dilution in 5% normal goat serum. Secondary antibody goat anti-mouse IgG Alexa Fluor 568 (Invitrogen, Cat. No. A-11004) was also applied at a 1:2000 dilution in 5% normal goat serum, along with Hoechst stain.

### Colocalization Analysis

For the colocalization analysis between internalized tau oligomers and the specified markers, we generated a mask for the non-tau channel by smoothing the image with a Gaussian kernel and applying the threshold_triangle function from the scikit-image library. The tau channel mask was created using the puncta identification method described in the section on lipid droplet quantification.

To quantify colocalization, the overlap coefficient was calculated as the ratio of the intersection area to the union area of the two masks. Additionally, Mander’s coefficient was determined by computing the mean intensity of the non-tau channel within the tau channel mask, normalized by the average intensity of the non-tau channel across the image.

### Tau Degradation Inhibitor Assay

Astrocytes were seeded at a density of 120,000 cells per well in a 24-well plate pre-coated with Matrigel and maintained in Astrocyte Media. After 24 hours, the media were replaced, and the cells were treated with 50 nM 2N4R-P301L tau oligomers conjugated to Alexa Fluor 647 (AF647) for 3 hours at 37°C. Following treatment, the cells were washed three times with PBS to remove any non-internalized tau. Fresh media containing 15 uM MG132 or 0.5 uM Bafilomycin A1 were added to the cells, which were then incubated for 16 hours. Cells were dissociated using Accutase, fixed with 4% paraformaldehyde at room temperature for 15 minutes, and stored in PBS at 4°C until flow cytometry analysis.

Flow cytometry was performed using a SONY MA900 cell sorter. For cell samples, gating was applied to exclude debris and isolate singlet populations, and the median AF647 fluorescence intensity of the singlet cells was measured to quantify intracellular tau levels.

### Tau Treatment RNA Sequencing

Differentiated astrocytes were plated in 60-mm dishes and cultured until they reached full confluency. Cells were treated with 50 uM of tau oligomers or vehicle (PBS) for 24 hours before extracting RNA. Four biological replicates were prepared for each condition and cell line. For RNA extraction, a protocol described in the RNA extraction section was followed.

Library preparation for RNA sequencing using NEBNext Ultra II Directional RNA Library Prep Kit (New England Biolabs, Cat. No. E7760S) with the NEBNext Poly(A) mRNA Magnetic Isolation Module (New England Biolabs, Cat. No. E7490S) and NEBNext Multiplex Oligos for Illumina (New England Biolabs, Cat. No. E7335S) and Next-Generation Sequencing were performed at the UCSB Biological Nanostructures Laboratory. Sequencing was done on an Illumina NextSeq 500 using a 75-cycle high-output kit.

### Tau Treatment Transcriptomic Analysis

RNA sequencing fastq files were quality-checked using FastQC and trimmed with Trimmomatic. The trimmed fastq files were then aligned to the GRCh38 human genome assembly with HISAT2. FeatureCounts was used to generate a count matrix by counting gene features. Low-expression genes were filtered by retaining rows with a minimum of 10 counts to ensure robust analysis.

Differential expression analysis was conducted with the DESeq2. The experimental design formula, “Genotype + Treatment + Genotype:Treatment”, was used to assess both independent and interactive effects of genotype and treatment. Genotype (WT or APOE3-Ch) and treatment (PBS or Tau-treated) were set as factors, and reference levels were specified to facilitate comparisons. The DESeq2 pipeline was executed, including normalization and dispersion estimation, followed by differential expression testing. Adjusted p-values (FDR) were used to identify significantly differentially expressed genes (DEGs), with a threshold of FDR ≤ 0.05.

Significant DEGs were further analyzed using GSEA to identify enriched biological pathways. Log2 fold change values of significant genes were ranked and mapped to the DEGs. Gene Ontology gene sets from the Molecular Signatures Database (MSigDB) were imported, and enrichment was performed using the fgsea package. Pathways with normalized enrichment scores (NES) and FDR ≤ 0.05 were identified as significantly enriched. Upregulated and downregulated pathways were visualized using horizontal bar plots, with pathways ranked by NES. Transparency was scaled to reflect FDR values.

Pathways of interest were selected based on GSEA results. Leading-edge genes contributing to pathway enrichment were extracted, and their expression profiles were visualized. Gene expression data were normalized (z-scored) across samples to highlight relative differences. Genes were grouped into treatment categories (WT untreated, WT treated, APOE3-Ch untreated, APOE3-Ch treated) to capture regulatory patterns.

### Statistical Analysis

All statistical analyses were conducted using R (v4.3.3), Python (v3.11.5), or GraphPad Prism 10. Details of the specific statistical methods are provided in the corresponding method sections. Data are presented as mean ± SD unless otherwise stated, with statistical significance defined as p < 0.05. All analysis parameters and scripts are available upon request to ensure transparency and reproducibility.

## Acknowledgments

We acknowledge support from the Center for Scientific Computing at the California Nanosystems Institute (CNSI, NSF grant CNS-1725797) for the availability of high-performance computing resources and support. This work used the Extreme Science and Engineering Discovery Environment, which is supported by the NSF grant ACI-1548562 (MCA05S027). We acknowledge the use of the NRI-MCDB Microscopy Facility and the Resonant Scanning Confocal supported by NSF MRI grant DBI-1625770. We acknowledge the use of the Biological Nanostructures Laboratory within the California NanoSystems Institute, supported by the University of California, Santa Barbara and the University of California, Office of the President. We acknowledge the use of the Mass Spectrometry Facility at University of Texas Medical Branch, supported by the Cancer Prevention Research Institute of Texas (CPRIT) awarded to Dr. William Russell (grant number: RP190682). Illustrations were created by Biorender.com.

## Author contributions

X.T. and K.S.K. designed research; X.T., E.K.R., A.A.N, I.H., J.A.S., Y.J.C, J.S., A.D., C.A., T.G. performed research; X.T. and K.S.K. analyzed data; and X.T. and K.S.K. wrote the paper.

## Data and materials availability

All data, code, and materials used in this study are available in the main text or the supplementary materials. Any additional information required to reproduce or extend the analyses presented here can be obtained from the corresponding authors upon request. Materials may be subject to transfer agreements.

---

**fig.S1:**
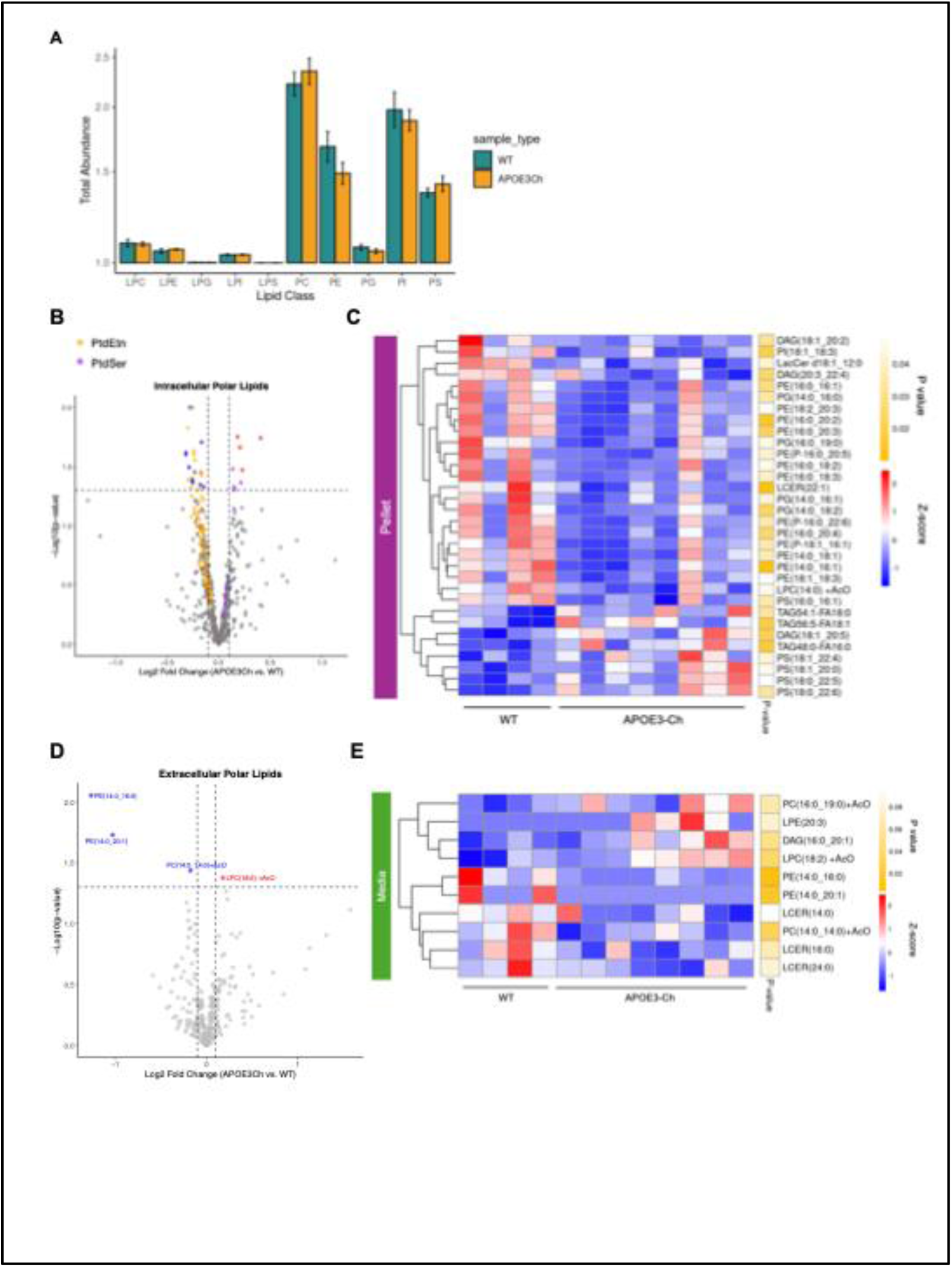
The polar lipid profiles of WT and APOE3Ch astrocytes. **(A)** Total lipid abundance (sum of normalized peak area) for each phospholipid class in the intracellular polar lipid panel, comparing APOE3-Ch and WT astrocytes. Error bars represent the SEM. **(B)** Volcano plot of intracellular polar lipids comparing APOE3-Ch and WT astrocytes with phosphatidylethanolamine (PtdEtn) and phosphatidylserine (PtdSer) classes highlighted in orange and purple, respectively. Lipids with significant upregulation (p-value < 0.05, log2 fold change > 0.1) are indicated by red points, and lipids with significant downregulation (p-value < 0.05, log2 fold change < 0.1) are indicated by blue points. **(C)** Heatmap showing differentially regulated lipids in the pellets (intracellular) of APOE3-Ch astrocytes compared to WT astrocytes. Lipid expression is represented as a z-score, calculated by subtracting the mean lipid quantity across all samples from the observed quantity and dividing by the standard deviation. Upregulated and downregulated lipids are color-coded. P-values were calculated using a linear model with empirical Bayes moderation of standard errors. **(D)** Volcano plot of extracellular polar lipids comparing APOE3-Ch and WT astrocytes. Lipids with significant upregulation (p-value < 0.05, log2 fold change > 0.1) are indicated by red points, and lipids with significant downregulation (p-value < 0.05, log2 fold change < 0.1) are indicated by blue points. **(E)** Heatmap showing differentially regulated lipids in the media (extracellular) of APOE3-Ch astrocytes compared to WT astrocytes. Lipid expression is represented as a z-score, calculated by subtracting the mean lipid quantity across all samples from the observed quantity and dividing by the standard deviation. Upregulated and downregulated lipids are color-coded. P-values were calculated using a linear model with empirical Bayes moderation of standard errors.

**fig.S2:**
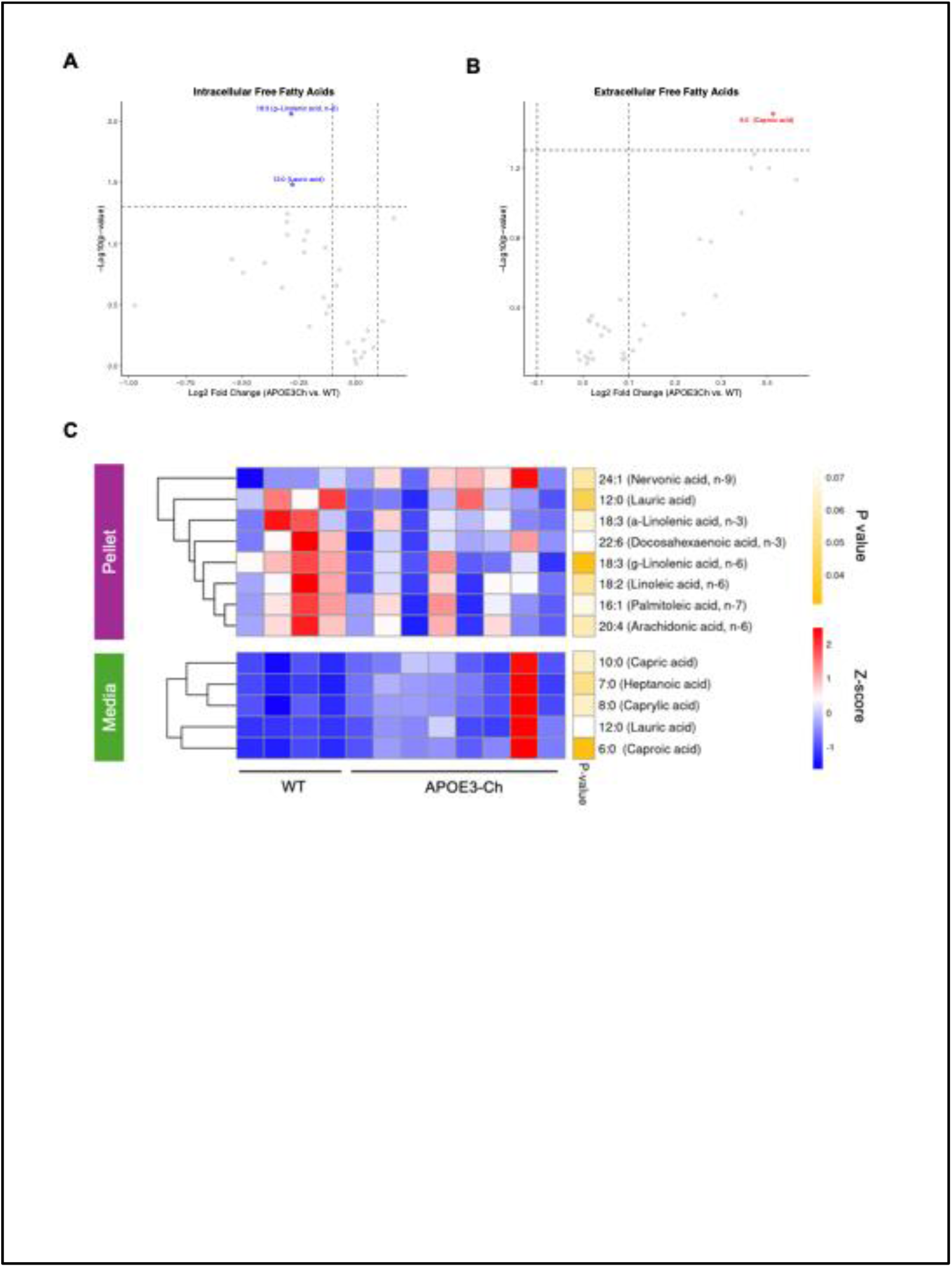
The free fatty acids (FFA) profiles of WT and APOE3Ch astrocytes. **(A)** Volcano plot of intracellular FFAs comparing APOE3-Ch and WT astrocytes. Lipids with significant upregulation (p-value < 0.05, log2 fold change > 0.1) are indicated by red points, and lipids with significant downregulation (p-value < 0.05, log2 fold change < 0.1) are indicated by blue points. **(B)** Volcano plot of extracellular FFAs comparing APOE3-Ch and WT astrocytes. Lipids with significant upregulation (p-value < 0.05, log2 fold change > 0.1) are indicated by red points, and lipids with significant downregulation (p-value < 0.05, log2 fold change < 0.1) are indicated by blue points. **(C)** Heatmap showing differentially regulated FFAs in the pellets (intracellular) and media (extracellular) of APOE3-Ch astrocytes compared to WT astrocytes. Lipid expression is represented as a z-score, calculated by subtracting the mean lipid quantity across all samples from the observed quantity and dividing by the standard deviation. Upregulated and downregulated lipids are color-coded. P-values were calculated using a linear model with empirical Bayes moderation of standard errors.

**fig.S3:**
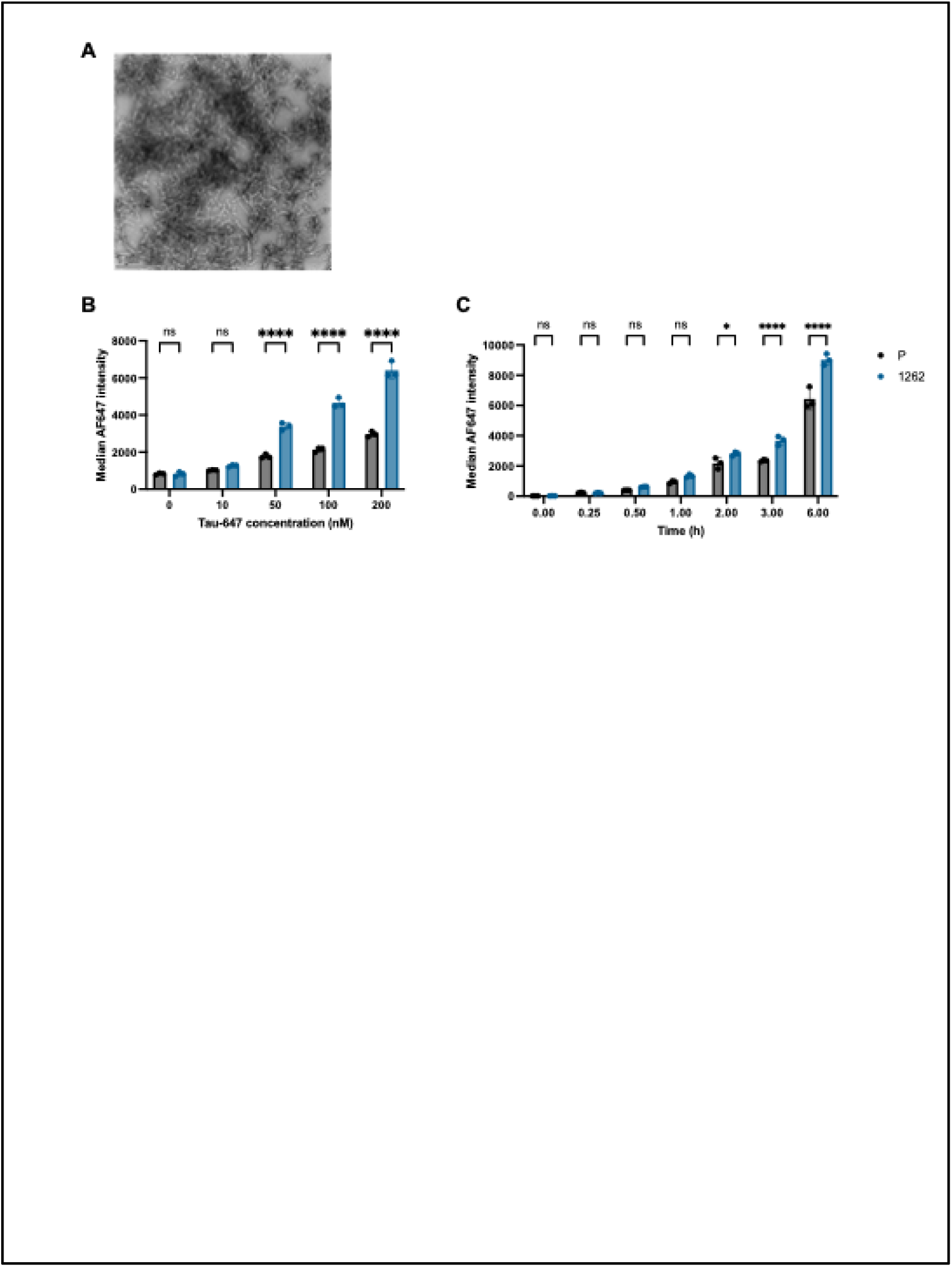
Concentration gradient and dynamics of tau uptake assay. **(A)** Representative TEM image of the 2N4R-P301L aggregates labeled with AF647 (Tau-AF647). **(B)** Measurement of the intracellular Tau-AF647 level in cells treated with different concentrations of Tau-AF647 for 3 hours (n=3, 10000 cells each). **(C)** Measurement of the intracellular Tau-AF647 level in cells treated with 50 nM of Tau-AF647 for various amounts of time (n=3, 10000 cells each). All data are expressed as mean ± s.d. with individual data points shown. Two-way ANOVA with Šídák’s multiple comparisons tests was performed to determine the significance unless otherwise specified. The results designated as “ns” are not significant; *p<0.05; **p<0.01; ***p<0.001;****p<0.0001.

**fig.S4:**
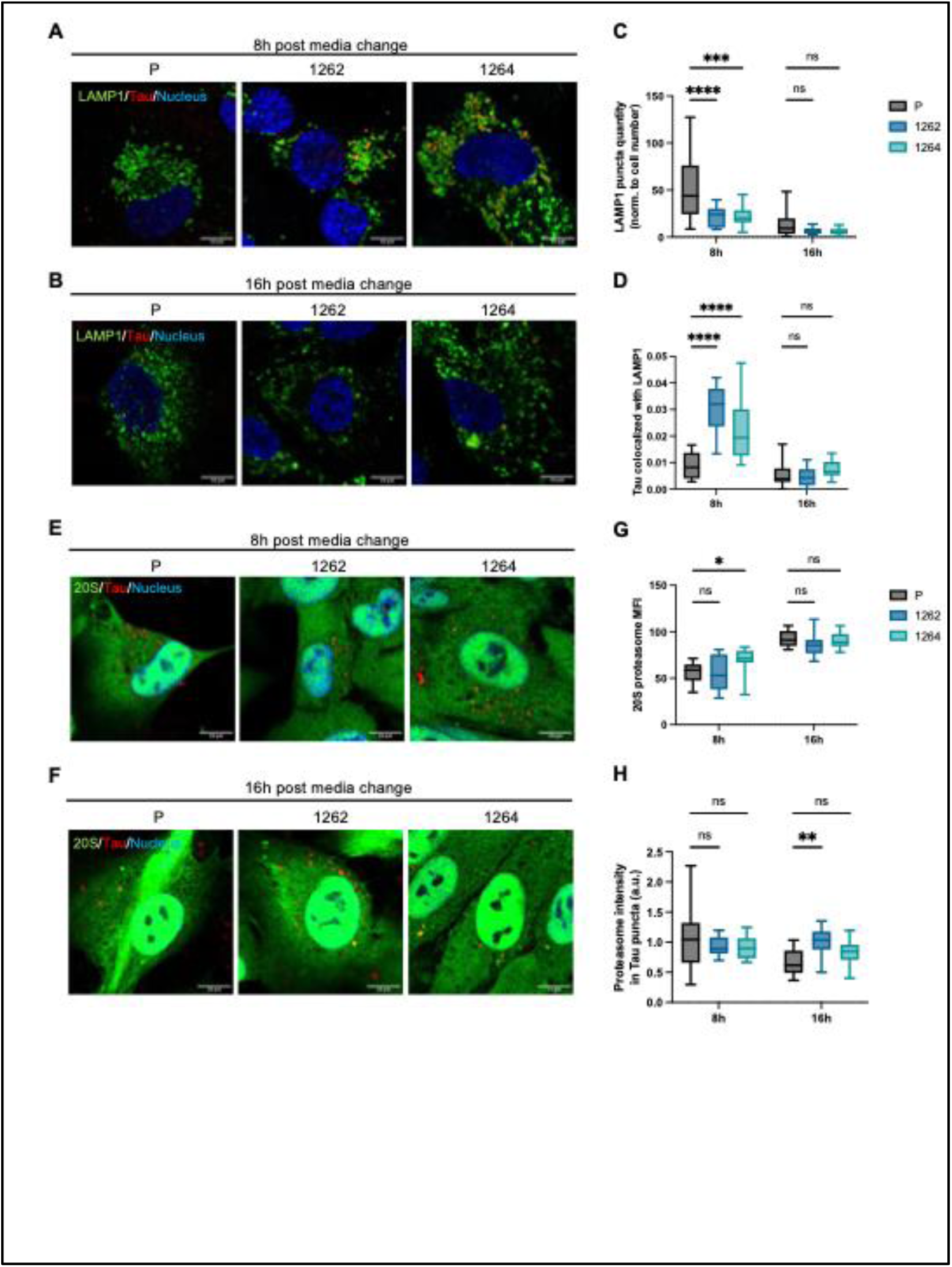
Exogenous Tau colocalizes with lysosome and proteasome. **(A)** Representative images of iPSC-derived astrocytes immunolabeled with anti-LAMP1, a lysosome marker, at 8 hours after media change. Green, LAMP1; red, Tau-AF647; blue, Hoechst 33342; scale bar = 10 µm **(B)** Representative images of iPSC-derived astrocytes immunolabeled with anti-LAMP1, a lysosome marker, at 16 hours after media change. Green, LAMP1; red, Tau-AF647; blue, Hoechst 33342; scale bar = 10 µm **(C)** Quantification of LAMP1 expression by the LAMP1+ puncta quantity normalized to the number of cells (n=12 FOVs taken from 4 biological replicates, 3 FOVs per biological replicate) **(D)** Quantification of Tau colocalized with LAMP1 (n=12 FOVs taken from 4 biological replicates, 3 FOVs per biological replicate) **(E)** Representative images of iPSC-derived astrocytes immunolabeled with anti-20S proteasome, at 8 hours after media change. Green, 20S proteasome; red, Tau-AF647; blue, Hoechst 33342; scale bar = 10 µm **(F)** Representative images of iPSC-derived astrocytes immunolabeled with anti-20S proteasome, at 16 hours after media change. Green, 20S proteasome; red, Tau-AF647; blue, Hoechst 33342; scale bar = 10 µm **(G)** Quantification of 20S proteasome expression by the mean fluorescence intensity (MFI) (n=12 FOVs taken from 4 biological replicates, 3 FOVs per biological replicate) **(H)** Quantification of the proteasome signal within Tau puncta (n=12 FOVs taken from 4 biological replicates, 3 FOVs per biological replicate) All data are expressed as mean ± s.d. with individual data points shown. Two-way ANOVA with uncorrected Fisher’s LSD tests was performed to determine the significance. The results designated as “ns” are not significant; *p<0.05; **p<0.01; ***p<0.001;****p<0.0001.

**fig.S5:**
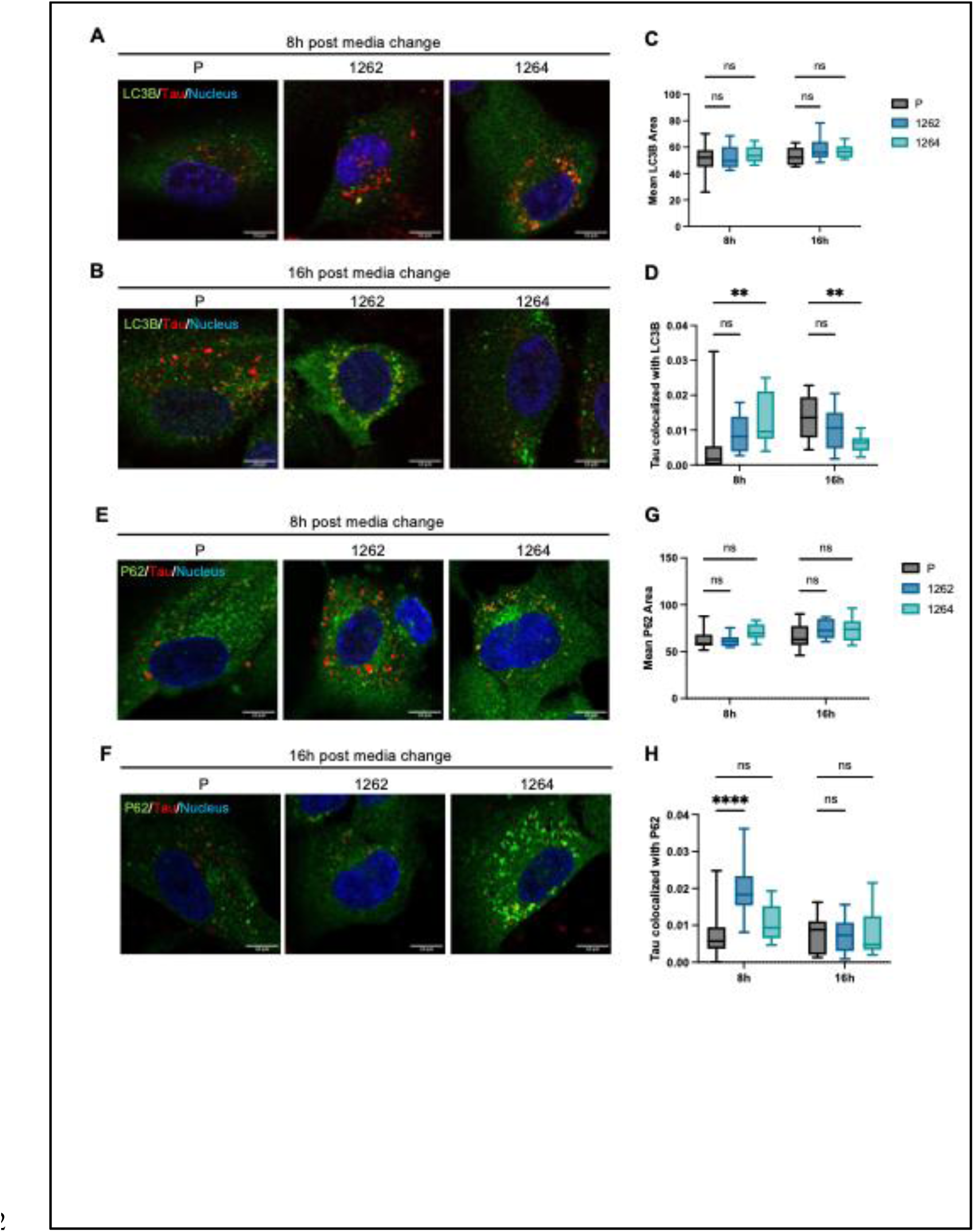
Exogenous Tau colocalizes with autophagosome and aggresome. **(A)** Representative images of iPSC-derived astrocytes immunolabeled with anti-LC3B, an autophagosome marker, at 8 hours after media change. Green, LC3B; red, Tau-AF647; blue, Hoechst 33342; scale bar = 10 µm **(B)** Representative images of iPSC-derived astrocytes immunolabeled with anti-LC3B, an autophagosome marker, at 16 hours after media change. Green, LC3B; red, Tau-AF647; blue, Hoechst 33342; scale bar = 10 µm **(C)** Quantification of LC3B expression by the mean LC3B area (n=12 FOVs taken from 4 biological replicates, 3 FOVs per biological replicate) **(D)** Quantification of Tau colocalized with LC3B (n=12 FOVs taken from 4 biological replicates, 3 FOVs per biological replicate) **(E)** Representative images of iPSC-derived astrocytes immunolabeled with anti-P62, at 8 hours after media change. Green, P62; red, Tau-AF647; blue, Hoechst 33342; scale bar = 10 µm **(F)** Representative images of iPSC-derived astrocytes immunolabeled with anti-P62, at 16 hours after media change. Green, P62; red, Tau-AF647; blue, Hoechst 33342; scale bar = 10 µm **(G)** Quantification of p62 expression by the mean p62 area (n=12 FOVs taken from 4 biological replicates, 3 FOVs per biological replicate) **(H)** Quantification of Tau colocalized with p62 (n=12 FOVs taken from 4 biological replicates, 3 FOVs per biological replicate) All data are expressed as mean ± s.d. with individual data points shown. Two-way ANOVA with uncorrected Fisher’s LSD tests was performed to determine the significance. The results designated as “ns” are not significant; *p<0.05; **p<0.01; ***p<0.001;****p<0.0001.

**fig.S6:**
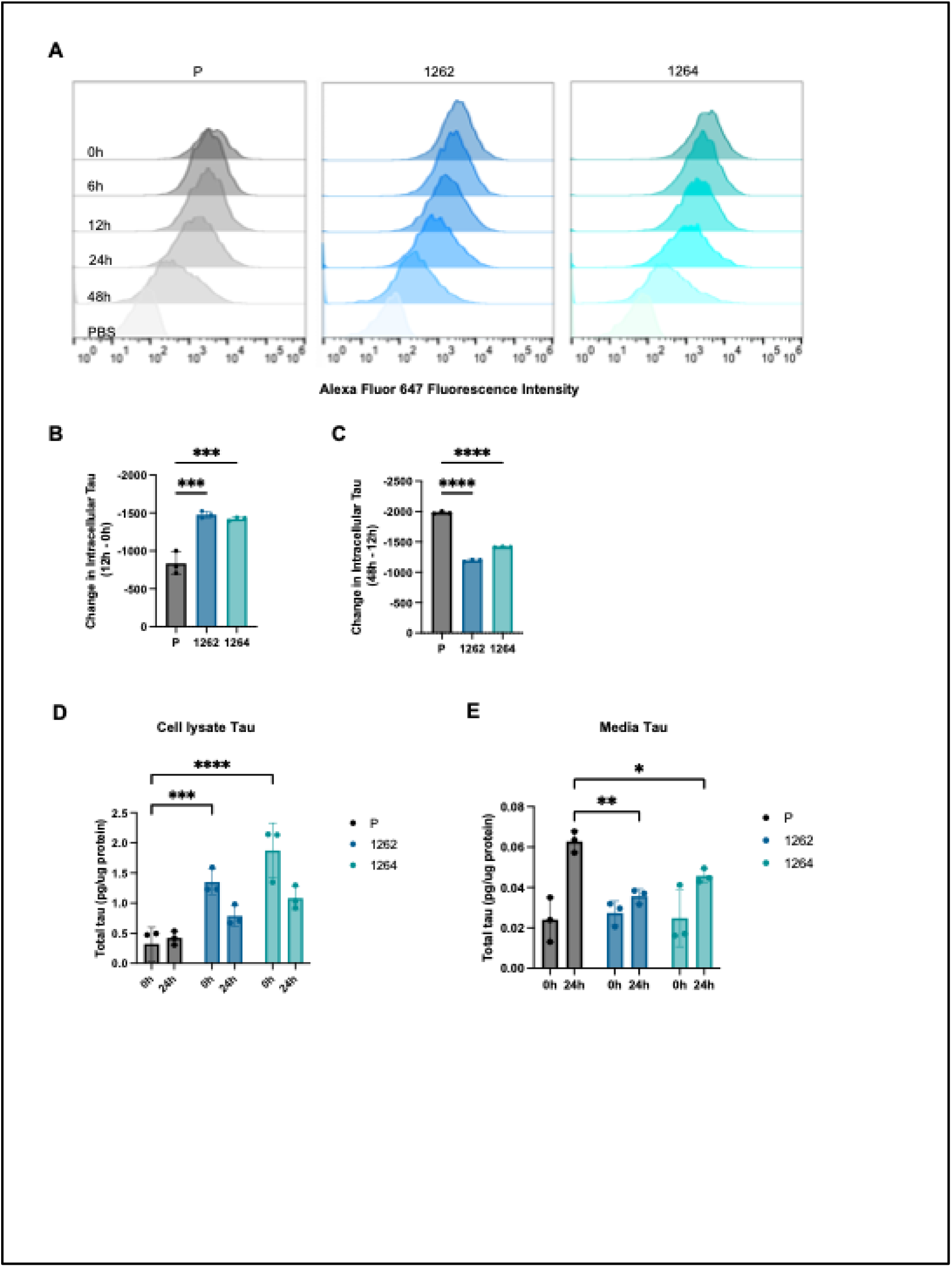
Dynamics of Tau Clearance in Astrocytes. **(A)** Representative overlaid histograms demonstrating the shift in fluorescence of cells treated with AF647 labeled tau (2N4R-P301L) aggregates (100nM, 3h) after media change at 0h, 6h, 12h, 24h, and 48h compared to cells treated with PBS. **(B)** Measurements of the change in intracellular Tau-AF647 level, comparing the median AF647 intensity of cells at 12 hours post media change to that of cells at 0 hour post media change. **(C)** Measurements of the change in intracellular Tau-AF647 level, comparing the median AF647 intensity of cells at 48 hours post media change to that of cells at 12 hours post media change. **(D)** Measurement of total tau concentration in the cell lysate with ELISA normalized to total protein concentration measured by BCA assays at 0 hours post media change and at 24 hours post media change **(E)** Measurement of total tau concentration in the media with ELISA normalized to total protein concentration measured by BCA assays at 0 hours post media change and at 24 hours post media change

